# Genomic analysis of the *Streptococcus anginosus* group; taxonomic implications, evolutionary insights and clinical considerations

**DOI:** 10.64898/2026.11.04.764143

**Authors:** Andrea Coleman, Joshua Liaw, Diane Maresco-Pennisi, Rylee Deehan, Patrick N.A. Harris, Anders Cervin, Oskar Edkins, Julian Zaugg, Seweryn Bialasiewicz

## Abstract

**Background:** The *Streptococcus anginosus* group (SAG) (*S. anginosus, S. intermedius,* and *S. constellatus*) causes serious, potentially life-threatening, invasive infections, often in otherwise healthy individuals. However, the group’s diversity is still poorly understood, and with it, the ability to identify prognostic markers of severe disease.

**Methods:** Long-read sequencing produced closed genomes for 29 isolates from 25 patients hospitalised primarily for invasive SAG infections, and 21 isolates from 20 non-hospitalised controls. Genomes were classified with GTDB-Tk and combined with 211 reference genomes. After dereplication, 207 SAG genomes (38 isolate, 169 reference) representing 206 strains were analysed using: GTDB classification, average nucleotide identity, multi-marker and core-genome SNP phylogenies, recombination-aware analyses, mobilome, CAZyme and gapseq metabolic profiling, and lineage-aware genome-wide association testing for invasive phenotype.

**Results:** Agreement between GTDB-Tk and MALDI-TOF taxonomic classifications was low (42.8%), including two *S. massiliensis* cases mis-identified by MALDI-TOF as SAG. *S. anginosus* resolved into four taxa (*S. anginosus*, *S. hominis*, *S. whileyi* and *S. vaginalis*). Phylogenies based on core genome single-nucleotide polymorphisms resolved these species into distinct clades, whereas 16S rRNA gene and multilocus sequence analysis phylogenies did not. The four ex-*S. anginosus* taxa showed a gradient of genomic cohesion, with *S. vaginalis* being near-clonal, in contrast to *S. anginosus,* which shared high levels of sequence ancestry with other neighbouring taxa. Carbohydrate utilisation was strongly lineage-associated, as was propensity for invasive disease. A ∼15Kbp glycosaminoglycan utilisation locus was found in the three taxa of highest disease severity propensity (*S. intermedius, S. constellatus, S. whileyi*) which predicts a direct host glycosaminoglycan feeding mechanism that may contribute to invasive disease. No single functional feature, gene or core-genome SNP was associated with invasiveness however after lineage-adjustment. Combined feature analysis suggests *S. vaginalis* has undergone a niche restriction and has evolved for a primary existence in the female urogenital tract.

**Conclusions:** Under-reported species may be mis-diagnosed as SAG invasive infections. SAG is more diverse than its current three-species definition, with the conventional *S. anginosus* species proposed to be classed into a complex of four distinct taxa. Invasive capacity appears to be an emergent system-level property rather than from individual features.

## 1 Introduction

*Streptococcus anginosus* Group (SAG) currently has three recognised species, *S. anginosus, S. constellatus,* and *S. intermedius* (1). Although SAG are commensal flora of the mouth, gastrointestinal, and genitourinary tracts, they also play a key role in invasive and pyogenic infections (2–4). The incidence of serious SAG-associated ear, nose and throat (ENT) infections in both children and adults has increased over time (4, 5). SAG are frequently detected in deep neck space infections, superficial suppurative lymphadenitis, and intracranial abscesses (6–9) and have become the most common *Streptococcus* species to cause sterile body site infections (4). In context of ENT, a case-series by The Texas’s Children’s Hospital documented 95 cases of SAG-complicated otitis media and sinusitis from 2011 to 2018, of which 100% required surgery, 37% required intensive care treatment and 8% had persistent neurological deficit, despite 76% of the patients being previously healthy (5). Although the increasing incidence is in part due to improved diagnostic methods, it is possible that strains are evolving with increasing virulence (4).

Previous sequence-based analyses suggest that the SAG is more diverse than the three recognised species. Jensen and colleges investigated 123 SAG isolates using multilocus sequencing analysis (MLSA) combined with 16S rRNA gene sequence and phenotypic characterisation (10). They identified seven distinct clusters, including three subspecies of *S. anginosus* (*S. anginosus* subspecies anginosus, *S. anginosus* genomosubspecies AJ1, *S. anginosus* subspecies *whileyi*)(10). Although SAG infections are treated clinically as a homogenous group, emerging evidence suggests that each species may have a distinct clinical phenotype. *S. intermedius* is strongly associated with purulent infections, with a propensity for the head, neck and central nervous system (3, 11, 12). *S. anginosus* and *S. constellatus* have been associated with bacteraemia, which carries a higher mortality rate of 10–29%, compared to SAG pyogenic infections (11–13). Accurate identification of species within SAG may enable more precise and complete epidemiological data and support prognostication. In the clinical setting, rapid identification using molecular methods may lead to earlier surgical intervention and rationalisation of antibiotic therapy supporting antimicrobial stewardship efforts.

Currently there is no gold standard for identification of SAG (3). Culture-based methods combined with biochemical commercial identification kits can provide accurate group-level identification but cannot distinguish individual species (3). Bacterial culture followed by identification with Matrix-Assisted Laser Desorption Ionization Time-of-Flight (MALDI-TOF) is showing increasingly accurate species-level identification, however, this method requires culture which can take several days, and relies on the acquisition of viable bacteria from clinical samples (14). Various PCR methods have been described, however, they are often poor at discriminating between *S. constellatus* and *S. intermedius,* and although they show promise, have yet to translate into clinical practice (3). Whole genome sequencing (WGS) is the most accurate means of identifying SAG species, enabling genome-level analysis for virulence factors, antibiotic resistance genes, as well as comparative genomics (15). Applied to isolates from the genitourinary tract, WGS has revealed a distinct “Urinae” group of *S. anginosus* which shows a primary tropism for the urinary tract (16, 17), as well as leading to the identification of genes essential for growth in an *S. anginosus* strain isolated from a bacteraemic patient (18). However, to our knowledge, WGS has not yet been applied to SAG isolates from invasive ENT infections.

Here we apply WGS to SAG isolates from ENT infections and compare them with isolates from healthy individuals and published SAG genomes. This exploratory genomic analysis aimed to clarify SAG taxonomy and investigate to its functional genomics, evolutionary history, and clinical implications. For this paper we have defined a ‘*Streptococcus anginosus* complex’(SAC) resulting from the division of the conventional *S. anginosus* species classification, which includes *S. anginosus*, *S. hominis*, *S. whileyi* and *S. vaginalis* (described in detail 3.2.2). Henceforth SAG will refer to SAC in addition to *S. constellatus* and *S. intermedius*).

## 2 Methods and Materials

A study protocol overview, including the analyses workflow, is illustrated in **Figure 1**.

**Figure 1.**
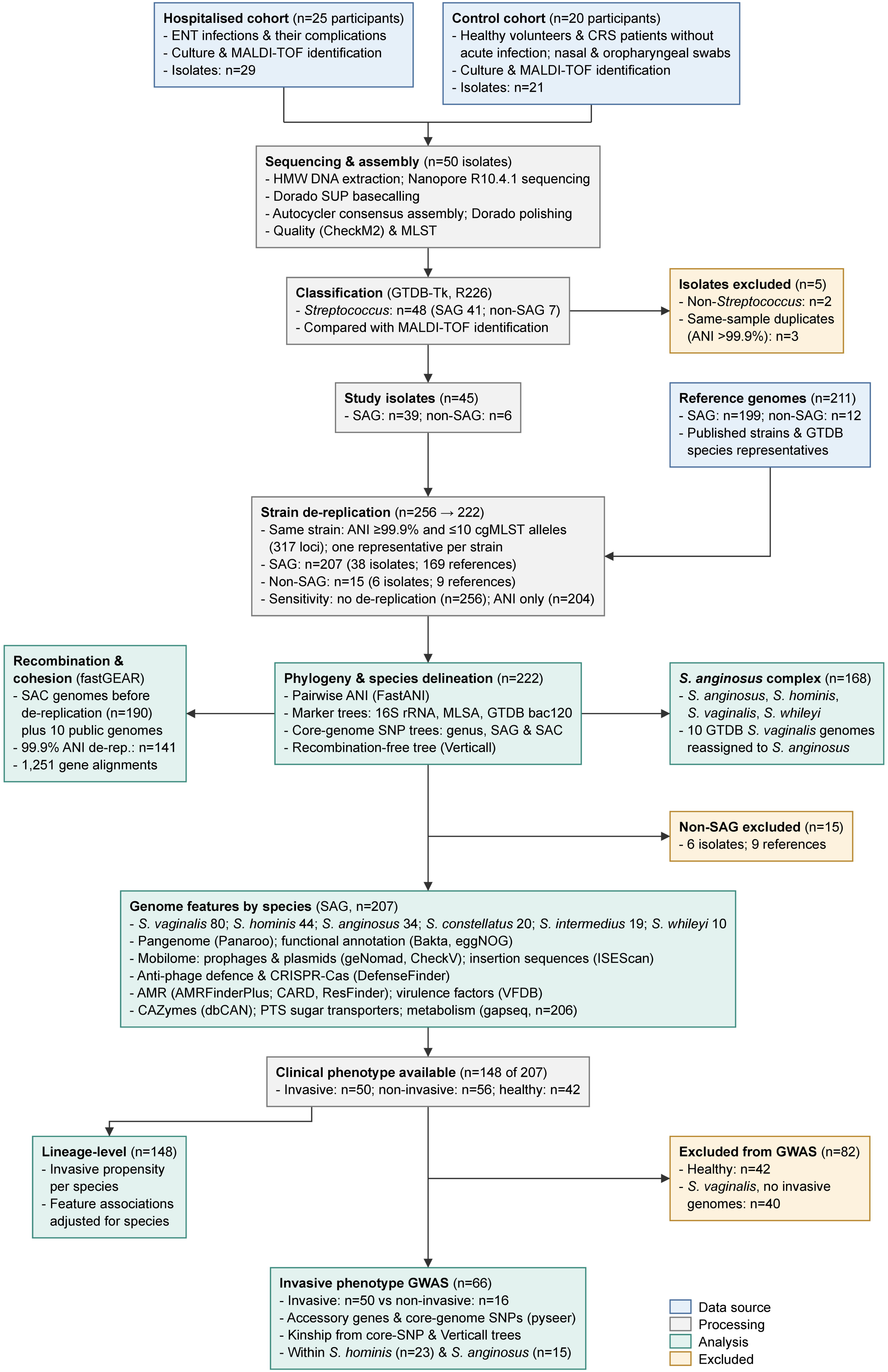
Study workflow overview. Isolates from 25 hospitalised patients and 20 control participants were sequenced and classified, combined with 211 reference genomes, and de-replicated to one genome per strain (222 genomes, 207 in the *S. anginosus* group). Box colours mark data sources, processing steps, analyses and exclusions. Abbreviations: ENT = Ear, Nose, Throat; CRS = Chronic Rhinosinusitis; HMW = High Molecular Weight; SUP = Super High Accuracy mode; GTDB-Tk = Genome Taxonomy Database Toolkit; MLSA = Multi-Locus Sequence Analysis; bac120 = GTDB 120 marker gene panel; ANI = Average Nucleotide Identity; GWAS = Genome Wide Association Study; SAG = *S. anginosus* group; SAC = *S. anginosus* complex; cgMLST = core-genome multi-locus sequence typing; MLST = multi-locus sequence typing; SNP = single-nucleotide polymorphism; AMR = antimicrobial resistance; VFDB = Virulence Factor Database; CAZymes = carbohydrate-active enzymes; PTS = phosphotransferase system; MALDI-TOF = matrix-assisted laser desorption/ionisation time-of-flight.

### 2.1 Study population

SAG isolates were collected prospectively from adults with ENT infections or their complications, including intracranial pus and bacteraemia, at a public tertiary hospital in Brisbane, Australia, between April and December 2024. All SAG isolates identified from ENT infections during this period were included, except from patients who did not consent. Bacteria were isolated as part of the routine diagnostic culture protocols, with MALDI-TOF (Vitek MS, BioMerieux) identification. Clinical data were obtained from patient medical records. Control isolates were obtained from an ongoing chronic rhinosinusitis (CRS) study, in which oropharyngeal and nasopharyngeal swabs from healthy adult volunteers or adults with CRS without acute infection were collected between March and August 2015. Swabs were cultured on selective media, and isolates were identified by MALDI-TOF as per routine diagnostic protocols. Ethics approval was obtained from Townsville Health and Hospital Service Human Research Ethics Committee (HREC/2022/QTHS/89471) for SAG-specific study collections and from the Royal Brisbane and Women’s Hospital Human Research Ethics Committee (HREC/14/QRBW/274) for the CRS-related collections.

### 2.2 Clinical definitions

Clinical outcomes were classified as healthy, invasive, or non-invasive according to predefined clinical criteria based on medical records of the SAG and CRS study participants, and for reference genomes, metadata and clinical information reported in the literature, where available (**Table 1**).

**Table 1.** Definitions of clinical outcome categories.

| **Category** | **Definition** |
|---|---|
| Healthy | Asymptomatic carriage (e.g. CRS controls), CRS without acute sinusitis, overactive bladder, and stress urinary incontinence |
| Non-invasive | Evidence of acute infection without complication. Examples include: acute sinusitis, dental abscess without deep neck involvement, urinary tract infections and cystitis. |
| Invasive | SAG infection beyond the typical anatomical site of commensal SAG colonization, including deep neck spaces, bacteraemia, endocarditis, intra-cranial sepsis and osteomyelitis. |

### 2.3 Isolate sequencing

Isolates were sub-cultured on HBA media to confirm purity, with a single colony incubated for 18– 25hrs in BHI broth culture at 37°C and 5% CO_2_. Horse blood supplementation was required in some instances. Genomic DNA from cell pellets was extracted with the Zymo Quick HMW DNA kit using the Cultured Bacterial and Fungal Cells protocol. Sequencing libraries were prepared using the native barcoding SQK-NBD114.24 kit as per manufacturer protocols (Oxford Nanopore Technologies, UK), with 24 barcoded and pooled samples being loaded onto R10.4.1 flow cells and run for 48 hours on a MinION MkD (Oxford Nanopore Technologies, UK).

### 2.4 Genome assembly, annotation and functional prediction

Sequence data was basecalled using Dorado (ver. 0.9.6+59ab908; model dna_r10.4.1_e8.2_400bps_sup@v5.0.0)(19) in super high accuracy mode, with adapters, primers and barcodes trimmed (--trim all). Any remaining adapters were identified and removed using Porechop (ver. 0.2.4; --discard_middle)(20). Reads were filtered with Filtlong (ver. 0.2.1; --min_length = 200, --min_mean_q = 9)(21). Filtered Nanopore reads were then *de novo* assembled with Autocycler (ver. 0.4.0) using the consensus of Canu (ver. 2.3)(22), Flye (ver. 2.9.5)(23), Hifiasm (ver. 0.25.0) (24–27), metaMDBG (ver. 1.1)(28), miniasm (ver. 0.3)(29), NECAT (ver. 0.0.1)(30), NextDenovo (ver. 2.5.2)(31), Plassembler (ver. 1.6.2)(32), and Raven (ver. 1.8.3)(33). The resulting consensus assemblies were then polished with Dorado polish. Assembled contigs were reorientated with dnaapler ‘all’ (ver. 1.0.1)(34), to start with one of dnaA (chromosomal), repA (plasmids) or terL (phage) genes (where applicable). Assembly completeness and contamination was evaluated using CheckM2 (ver. 1.0.2)(35), and taxonomy assigned to each using the Genome Taxonomy Database Toolkit (GTDB-Tk; ver. 2.4.1; with reference to GTDB R10-RS226)(36–38). Protein-coding sequences were predicted using Pyrodigal (version 3.6.3)(39) as part of Bakta (version 1.11.0)(40), and functionally annotated. Functional annotations were also generated using eggNOG Mapper (ver. 2.1.15; --sensmode sensitive, -m diamond) (41).

Antimicrobial resistance (AMR) and virulence genes were identified with AMRFinderPlus (ver. 4.0.22)(42) and ABRicate (ver. 1.4.0, https://github.com/tseemann/ABRICATE; --minid 60, -- mincov 70, --db card/vfdb/resfinder)(43–45). Insertion sequence elements were identified with ISEScan (ver. 1.7.2.3)(46). Carbohydrate-acting enzymes (CAZymes) were annotated with dbCAN (ver. 5.2.9) (47), with family membership taken as the union of calls from its three constituent search tools (HMMER, dbCAN_sub, DIAMOND). Antiphage defence systems were predicted with DefenseFinder (ver. 3.0.0) (48–50).

Read and assembly stats were calculated with SeqKit (ver. 2.8.1)(51). Finally, for each assembly, the fraction covered by at least one read, the mean coverage and the number of reads mapping were calculated by first mapping the respective reads with minimap2 (ver. 2.28-r1209) as part of CoverM (ver. 0.7.0)(52), and then removing low quality read mappings with CoverM ‘filter’ (minimum identity of 90% and minimum aligned length of 75%).

### 2.5 Reference genomes

A total of 199 SAG and 12 other study-relevant reference *Streptococcus* genomes were included in the present study for comparative purposes, which were identified from both clinically relevant strains described in the literature and species representatives from the Genome Taxonomy Database (GTDB)(36, 53, 54) (**Supplementary Table 1**). Representative genomes were chosen to complement the lineages identified among the sequenced isolates, and all 211 were processed using the same methods as the isolates. All but one were of high quality according to CheckM2 (≥ 90% completeness, ≤ 5% contamination). The exception, a medium-quality *S. constellatus* assembly (GCA_000223295.2; 88.6% completeness, 3.0% contamination), was retained for all analyses except metabolic prediction. Pairwise average nucleotide identity (ANI) between all isolate and reference genomes was calculated using FastANI (ver. 1.33)(55).

### 2.6 Genome dataset curation and strain-level dereplication

Genomes were dereplicated to one representative per strain before comparative analyses, so that repeat sequencing of the same strain was not counted as independent observations. Strains were defined using both ANI and core-genome multilocus sequence typing (cgMLST) allelic distance. For cgMLST, a study-specific whole-genome MLST (wgMLST) schema was created from all isolate and reference genomes with the chewBBACA (ver. 3.3.10)(56) ‘CreateSchema’ module, using a Pyrodigal training file built from the *S. anginosus* type strain NCTC 10713. Alleles were called using ‘AlleleCall’, paralogous loci removed with ‘RemoveGenes’, and ‘ExtractCgMLST’ was used to define a strict core of 317 loci called in every genome. Pairwise allelic distances were calculated with cgmlst-dists (ver. 0.4.0; https://github.com/tseemann/cgmlst-dists).

Two genomes were considered the same strain only if they shared ≥ 99.9% ANI (FastANI; mean of the two reciprocal estimates) and differed by ≤ 10 cgMLST alleles. ANI distance (100 − ANI) and allelic distance were each clustered by complete-linkage hierarchical clustering (R hclust) and cut at the corresponding threshold, so that every pair of genomes within a cluster met that threshold. Strains were defined as genomes that clustered together by both measures. The allelic threshold was calibrated on multiple independently sequenced reference genomes of the *S. anginosus* type strain, which differed by at most 7 alleles.

One representative genome was retained per strain, prioritising genomes with a clinical phenotype, then those with the fewest contigs, highest CheckM2 completeness, lowest CheckM2 contamination, and highest N50. Where genomes of the same strain had different clinical phenotypes, one representative was retained per phenotype, as these represent recovery of the same strain from individuals with different clinical phenotypes rather than redundant sampling. All downstream comparative analyses used the dereplicated set (with exception of fastGEAR, see below).

### 2.7 Phylogenetic analysis

To investigate the phylogenetic relationships among the detected *Streptococcus* species in detail, we generated multiple phylogenetic trees based on the alignment of different genomic markers: i) full 16S rRNA genes, ii) multilocus sequence analysis (MLSA)(57) of seven house-keeping genes, *iii)* the 120-gene single-copy marker genes used by GTDB, and *iv*) core genome single-nucleotide polymorphisms (SNPs).

For the 16S tree, Bakta-annotated 16S rRNA gene sequences with a minimum length of 1,300 bp were aligned with SSU-ALIGN (ver. 0.1.1)(58). Seven Bakta-annotated house-keeping genes (*map*, *pfl*, *pyk*, *ppaC*, *rpoB*, *soda*, and *tuf)* were used for the MLSA tree, with only genomes containing all seven genes being retained for further analysis. Amino acid sequences translated for each gene were aligned separately with MAFFT (ver. 7.455; --localpair) and concatenated. Similarly, the GTDB bac120 marker genes were translated and the amino acid sequences concatenated and aligned. Prior to tree generation, all alignments underwent trimming with trimAl (ver. 1.4.1; -automated1) to remove poorly aligned regions. RAxML was used to generate all maximum-likelihood trees, with the general time-reversible nucleotide substitution model with gamma correction for site variation (GTRGAMMA) model being applied to the 16S rRNA gene alignment, and the PROTCATLG model being applied to the MLSA and GTDB bac120 alignments. Number of bootstraps were determined by the autoMRE criterion (59).

A pangenome was constructed separately with Panaroo (ver. 1.8.0; --clean-mode strict, -- core_threshold 0.95, --threshold 0.90, --family_threshold 0.7, --len_dif_percent 0.95)(60) for SAC (*S. anginosus*, *S. hominis*, *S. vaginalis*, and *S. whileyi*), SAG as redefined here (SAC + S. *intermedius* and *S. constellatus*), and all *Streptococcus* (SAG plus the other *Streptococcus* genomes in the dataset). A SNP-only alignment was then produced from the resulting filtered core gene alignment with SNP-sites (ver. 2.5.1)(61). A maximum-likelihood phylogenetic tree was then constructed for the core gene SNP alignments using RAxML (ver. 8.2.12)(62) using GTRGAMMA and bootstraps determined by the autoMRE criterion.

Finally, to account for potential recombination among the genomes, we also constructed a recombination-corrected phylogeny with Verticall (ver. 0.4.3) using the distance tree workflow (63) and FastME (ver. 2.1.6.3) (64). All phylogenetic trees were visualised using tvBOT (65).

### 2.8 Virus and plasmid prediction

Viral sequences were identified and extracted using geNomad (ver. 1.11.0)(66) using conservative parameter settings (n_hallmarks >= 1, virus_score >= 0.8, marker_enrichment >= 1.5) with modifications to increase stringency (virus/plasmid score ≥90, and length ≥ 1,000bp). Host flanking regions for predicted viral sequences were trimmed using CheckV (ver. 1.0.3) (67). Viral and plasmid sequences were clustered based on pairwise ANI using the script aniclust.py within CheckV; Clustering was performed on sequences sharing 85% coverage, at both 70% and 95% identity thresholds (--min_ani 70 or 95, --min_tcov 85, --min_qcov 0). Only prophages assessed by CheckV as high-quality or complete were retained for analysis.

### 2.9 fastGEAR taxon cohesion and recombination

Recombination within the SAC was inferred with fastGEAR (68) (MATLAB Runtime 9.0.1, R2016a). The analysis used a separate set of 200 genomes, comprising those from the main dataset before dereplication, excluding two (one with an incongruent GTDB-Tk species assignment, one suppressed by NCBI), together with ten additional public *S. anginosus* and *S. whileyi* genomes included to increase resolution for these taxa (**Supplementary Table 1**). Gene alignments for fastGEAR were generated from a pangenome constructed with Panaroo (ver. 1.8.0; --clean-mode moderate, --threshold 0.95, --family_threshold 0.7, --len_dif_percent 0.95), with nucleotide alignments of all gene clusters generated with MAFFT(69) (-a pan).

To prevent near-identical genomes from inflating recombination counts and estimates of taxon cohesion, genomes were dereplicated by complete-linkage clustering of pairwise ANI at 99.9% as estimated by skani (ver. 0.3.2) (70). The representative of each cluster was the member present in the most gene alignments of an initial fastGEAR run on all 200 genomes. ANI alone was used because, for recombination inference, near-identical genomes are redundant whether they derive from a single culture or from different patients. fastGEAR was then rerun on the alignments of 1,251 gene clusters reduced to the resulting 141 genomes (*S. anginosus* 33, *S. hominis* 39, *S. vaginalis* 61, *S. whileyi* 8). Following Mostowy et al. (68), recombinations present in a subset of the genomes of a fastGEAR lineage were classed as recent, and those present in all genomes of a lineage as ancestral, having occurred before the lineage’s most recent common ancestor. Recent recombination segments that overlapped in position, came from the same donor lineage and occurred in genomes of the same recipient lineage were grouped into one event, reproducing fastGEAR’s own event count for every gene. Events reaching genomes of several taxa were counted once per recipient taxon, as a single transfer into a shared ancestor cannot be distinguished from separate transfers. As the direction of ancestral recombinations is difficult to determine (68), these were analysed as unordered pairs of lineages.

A segment’s donor was assigned to a taxon when at least 90% of the genomes in its donor lineage belonged to that taxon and was otherwise classed as mixed. Taxon cohesion, the mean proportion of shared ancestry between pairs of genomes within a taxon after accounting for recombination (68), was tested against 9,999 permutations of taxon labels. Per-genome recombination rates were compared between taxa with Kruskal–Wallis and pairwise Wilcoxon rank-sum tests with Benjamini– Hochberg correction. To distinguish recent transfer from retained ancestral variation, the similarity of each between-taxon segment to the closest genome of its donor lineage was compared, over the same alignment columns, with that of unrecombined recipient-taxon alleles.

### 2.10 Invasive phenotype GWAS

Associations between gene presence/absence and invasiveness were tested with a linear mixed model in pyseer (ver. 1.4.3)(71). Analyses were run separately on the SAC, SAG and all *Streptococcus* genome sets, each with its own pangenome, and repeated without *S. vaginalis*, which contained no invasive genomes. Population structure was accounted for with kinship matrices derived from the core-genome SNP (RAxML) and recombination-corrected (Verticall) trees, in separate analyses, to test whether results depended on recombination distorting the core-genome tree. Genes present in 15–85% of genomes were tested, and significance was assessed by Bonferroni correction over the number of unique presence/absence patterns across the analysed genomes. Heritability (h²) and genomic inflation (λ) were reported for each analysis, indicating respectively how much of the phenotype tracked phylogenetic relatedness and p-values were well calibrated.

The same tests were repeated on core-genome SNPs with the same trees and frequency filter, after decomposing multiallelic sites with bcftools (ver. 1.24; 9–35% of sites). Species-level effects were tested by comparing the proportion of invasive genomes in each species with all other species (Fisher’s exact test), Bonferroni-corrected for the number of species tested. To test for associations independent of species identity, accessory genes and core-genome SNPs were also tested within *S. hominis* and *S. anginosus*, the only species with both phenotypes represented by more than one genome, using species-specific kinship matrices.

### 2.11 Insertion sequence and prophage/plasmid abundance and composition

Insertion sequence copy number was compared between species with negative-binomial generalised linear models offset by log CDS (MASS ver. 7.3-65), using likelihood-ratio tests with Benjamini-Hochberg-corrected pairwise contrasts (emmeans ver. 2.0.4). Pairwise contrasts were limited to species with at least three genomes. IS family composition was compared by permutational multivariate analysis of variance (PERMANOVA) (72) on Bray-Curtis dissimilarities of per-genome family densities (copies per 1,000 CDS), and homogeneity of multivariate dispersions by permutation tests (PERMDISP) (73), both in vegan (ver. 2.7-5; 999 permutations)(74). *S. vaginalis* (80 of 207 SAG genomes, all of them reference genomes and none invasive) confounds species with symptom group and was excluded throughout as a sensitivity analysis.

Prophage and plasmid prevalence between species were tested by Fisher’s exact test (each species against the rest, Benjamini-Hochberg corrected). Copy number per genome was compared between species by negative-binomial generalised linear models offset by log CDS, with Benjamini– Hochberg-corrected pairwise contrasts

### 2.12 Functional flexibility (gene-content richness vs. clinical severity)

Functional flexibility was assessed with seven gene-content metrics, each calculated per genome as richness (the number of distinct types present) and abundance (the number of copies): virulence-factor categories (VFDB, abricate), AMR genes (CARD and ResFinder, abricate), defence-system types (DefenseFinder), IS families (ISEScan), prophage clusters (geNomad; CheckV high-quality or complete, clustered at 70% ANI), CAZyme families (dbCAN), and PTS sugar-transport substrates (Bakta product annotations matched to a fixed list of 18 substrates). Richness and abundance were normalised to counts per 1,000 CDS, with a pseudocount of one. Severity propensity per species was defined as the proportion of its phenotyped genomes that were invasive (Wilson score 95% CI) and was correlated with each metric by Spearman rank across species with at least five phenotyped genomes. Correlations were evaluated through leave-one-species-out refitting, exclusion of *S. vaginalis*, and restriction to sequenced isolates. The CAZyme correlation was also recalculated counting only families called by at least two of dbCAN’s three tools. CRISPR-Cas presence and subtype (DefenseFinder) were examined separately, as the proportion of genomes per species carrying a Cas system (Wilson score 95% CI), alongside severity propensity.

### 2.13 Metabolic capability prediction

Metabolic capability was inferred from Bakta-predicted proteins with gapseq (ver. 2.1.0; sequence database ver. 1.5, Bacteria; Gram-positive setting)(75), using only the homology-based ‘find’ and ‘find-transport’ modules, without gap-filling. One genome (GCA_000223295.2, no clinical phenotype) was excluded because its assembly carried extensive frameshift errors. Reactions were scored present only when supported by a homology hit (gapseq status ‘good_blast’), and pathways when at least 80% of their reactions were detected, or at least 66% including all key reactions. Each predicted transporter was counted in two ways: by the compound it transports (what can be taken up, identified by ModelSEED compound ID(76)) and by its Transporter Classification family (the type of transport system).

Jaccard distances between genomes were calculated on binary feature presence/absence and compared between species by PERMANOVA (999 permutations). To test whether metabolic content tracks shared ancestry, the Jaccard distance on reaction presence/absence was correlated with cophenetic distance from the Verticall tree using Mantel tests (Spearman, 999 permutations), across all genomes and within each species. Species-diagnostic features were identified by pairwise Fisher’s exact tests between species (Benjamini-Hochberg corrected). Features were retained at adjusted p < 0.05 with a prevalence difference of at least 80 percentage points between the two species (e.g. 90% vs 10%). Features were also assessed independently by a pyseer linear mixed model of each species against the rest, using the Verticall-derived kinship matrix.

Association with the invasive phenotype was tested in pyseer, as described for the accessory gene analysis, with kinship matrices derived from the core-genome SNP and Verticall trees in separate analyses. Reactions and transported compounds (by gapseq compound name) that varied between genomes were collapsed into blocks of identical presence/absence pattern, so that features always inherited together, such as the reactions of a single pathway, were counted once. Block composition was compared by PERMANOVA (Jaccard distance, 999 unrestricted permutations) across genomes with a clinical phenotype, testing invasive status (invasive vs healthy and non-invasive combined) after accounting for taxon.

### 2.14 Data processing and other analyses

All other data processing and analyses were performed, and visualisations generated, in R (ver. 4.6.1). Ordination plots based on accessory gene presence/absence were created with ggplot2 (ver. 4.0.3)(77), ggnewscale (ver. 0.5.2) and vegan (2.7-5)(74), and heatmaps with ComplexHeatmap (ver. 2.28.0)(78). Some analyses were conducted on the Galaxy Australia platform(79).

## 3 Results

### 3.1 Sampling and processing

#### 3.1.1 Hospitalised study participant isolates

Twenty-five participants were included, 22 with invasive infections (median age= 57, interquartile range (IQR)= 37-68) and three with non-invasive infections (median age= 62, IQR= 42-70) (**Table 2**). Two-thirds the participants had deep neck space infections (n=14, 64%) and 60% (n=15) experienced serious complications due to their SAG infection (**Table 2, Supplementary Table 2).** Most (76%, n=19) patients had no clinically significant co-morbidities **(Supplementary Table 2)**. In total, 29 isolates were obtained (*S. anginosus* n=10, *S. constellatus* n=10, *S. intermedius* n=7, and “S. milleri group” n=2); all but two were identified to the species-level using MALDI-TOF **(Supplementary Table 2).** Five (23%) participants had polymicrobial infections. In four participants, two isolates from the same sampling were obtained; in each case the paired isolates had matching MALDI-TOF identifications.

**Table 2:**
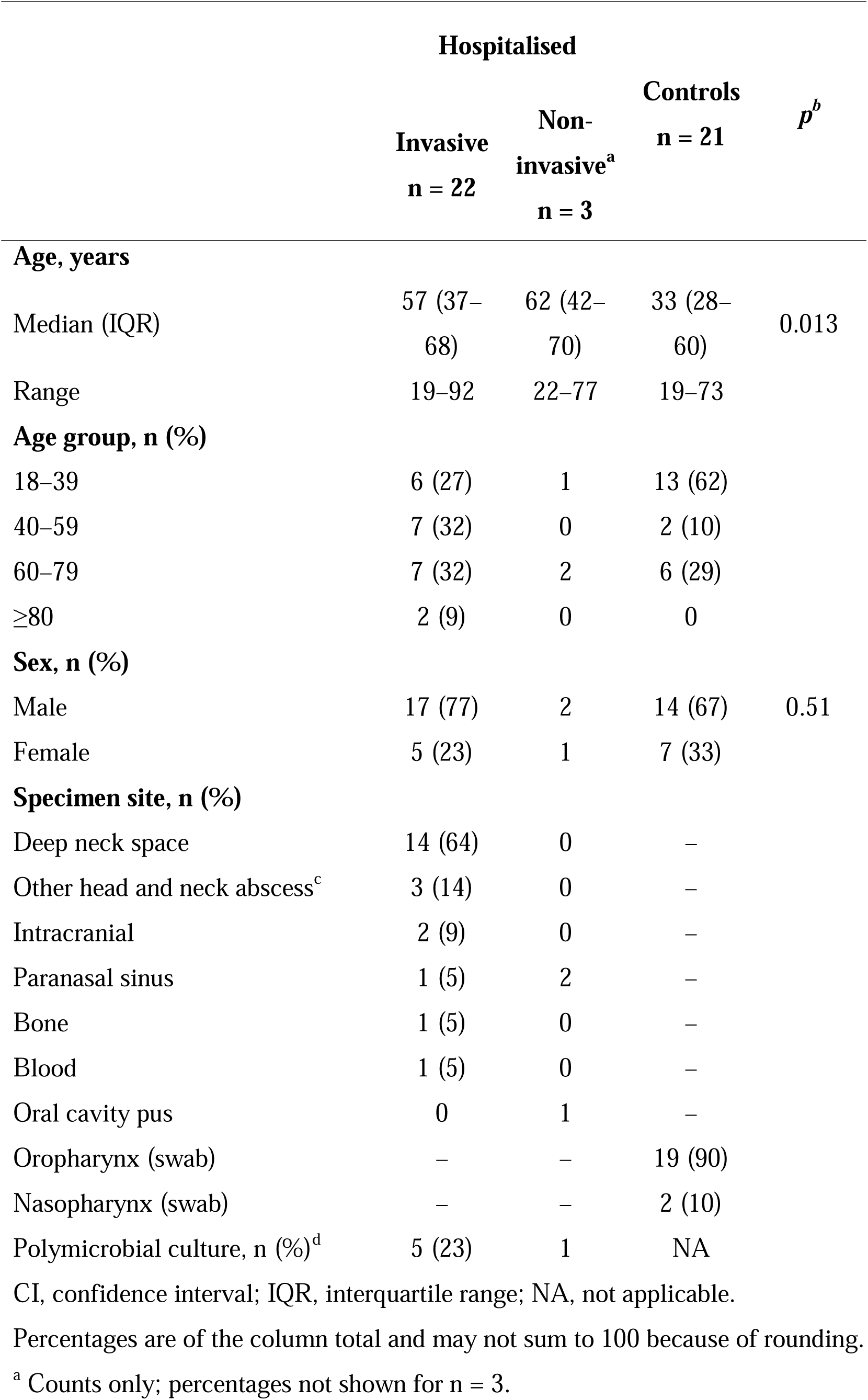
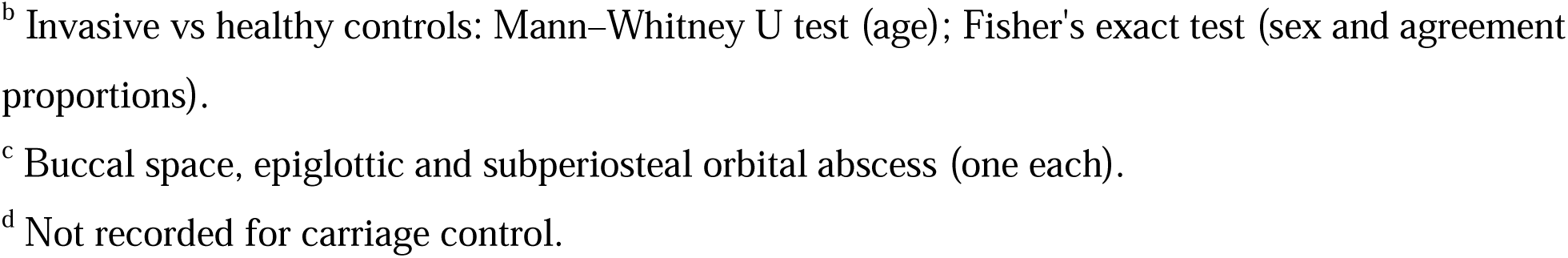
Summary of demographic and clinical information of participants and healthy controls.

|  | **Hospitalised** |  | **Controls<br>n = 21** | ***p***<sup>***b***</sup> |
|---|---|---|---|---|
|  | **Invasive<br>n = 22** | **Non-invasive<sup>a</sup><br>n = 3** |  |  |
| **Age, years** |  |  |  |  |
| Median (IQR) | 57 (37–68) | 62 (42–70) | 33 (28–60) | 0.013 |
| Range | 19–92 | 22–77 | 19–73 |  |
| **Age group, n (%)** |  |  |  |  |
| 18–39 | 6 (27) | 1 | 13 (62) |  |
| 40–59 | 7 (32) | 0 | 2 (10) |  |
| 60–79 | 7 (32) | 2 | 6 (29) |  |
| ≥80 | 2 (9) | 0 | 0 |  |
| **Sex, n (%)** |  |  |  |  |
| Male | 17 (77) | 2 | 14 (67) | 0.51 |
| Female | 5 (23) | 1 | 7 (33) |  |
| **Specimen site, n (%)** |  |  |  |  |
| Deep neck space | 14 (64) | 0 | – |  |
| Other head and neck abscess<sup>c</sup> | 3 (14) | 0 | – |  |
| Intracranial | 2 (9) | 0 | – |  |
| Paranasal sinus | 1 (5) | 2 | – |  |
| Bone | 1 (5) | 0 | – |  |
| Blood | 1 (5) | 0 | – |  |
| Oral cavity pus | 0 | 1 | – |  |
| Oropharynx (swab) | – | – | 19 (90) |  |
| Nasopharynx (swab) | – | – | 2 (10) |  |
| Polymicrobial culture, n (%)<sup>d</sup> | 5 (23) | 1 | NA |  |

CI, confidence interval; IQR, interquartile range; NA, not applicable.
Percentages are of the column total and may not sum to 100 because of rounding.
<sup>a</sup> Counts only; percentages not shown for n = 3.
<sup>b</sup> Invasive vs healthy controls: Mann–Whitney U test (age); Fisher's exact test (sex and agreement proportions).
<sup>c</sup> Buccal space, epiglottic and subperiosteal orbital abscess (one each).
<sup>d</sup> Not recorded for carriage control.

#### 3.1.2 Control isolates

Twenty-one SAG isolates (*S. anginosus* n=10, *S. constellatus* n=8, and *S. intermedius* n=3) from 20 CRS study participants (median age 33, IQR=19-73) were sequenced, including two *S. constellatus* isolates obtained from the same subject swab. All subjects were classified as “healthy” for the purposes of the comparative phenotype analyses (**Table 2**, **Supplementary Table 2**).

#### 3.1.3 Isolate sequencing

Across three flow cells, 38.96Gbp of sequence data were generated, with mean outputs of 215,145 reads, 700,068,336bp, 9,617 N50, and average read Q-scores of 17.57 per isolate (**Supplementary Table 3**). All isolate genomes were assembled into singular, closed contigs. Assembled genomes had a mean coverage of 325x/genome, with mean 99.99% completeness and 0.21% contamination scores.

### 3.2 Taxonomic classification

#### 3.2.1 Genomic classification is discordant with MALDI-TOF

Based on GTDB taxonomy, 14 bacterial species were characterised, resulting in a 42% (21/50) isolate agreement with the original MALDI-TOF identifications (**Supplementary Table 2**). The majority of discordant isolates (44.8%, 13/29) were disagreements within a *S. anginosus* complex (SAC), including the two isolates originally unable to be resolved to the species level, followed by intra-SAG (24.1%, 7/29), inter-Streptococcus group (24.1%, 7/29), and trans-Phylum (6.9%, 2/29) disagreements. Of note, three of the hospitalised cohort isolates (two from the same subject) resolved to *S. massiliensis* by GTDB and were misidentified by MALDI-TOF as either *S. anginosus* (U9494 and U8768) or *S. constellatus* (U4991) despite having near-identical genomes (ANI >99.9%). The three isolates came from two infections that were oral in origin and classified as invasive (**Supplementary Table 2**). Finally, two of isolates were identified as belonging to different Phyla (**Supplementary Table 2**); these were considered to represent either human handling errors or contamination and were also excluded from further analyses.

#### 3.2.2 ANI clustering and high-resolution phylogeny support *Streptococcus* classification

Pairwise genome ANI analyses of combined study and reference genomes (n=222) showed that *Streptococcus* isolates outside of the SAG formed distinct clusters corresponding to either *S. massiliensis* or *Streptococcus mitis* group (comprising *S. mitis*_BM, *S.* sp030546305*, S.* sp013277555, and an uncharacterised *Streptococcus*). While the *S. constellatus* and *S. intermedius* members of the SAG also formed defined clusters, *S. anginosus* formed a larger, more diverse cluster with dense sub-populations (**Figure 2**, **Supplementary Figure 1**). These corresponded to *S. anginosus* subspecies *whileyi* (10), *S. hominis* (80)*, S. anginosus*, and the *S. anginosus* “Group 2”/”Urinae” (16, 17), which is classified as *S. vaginalis* (81). The core-genome SNP phylogeny also supported the division of *S. anginosus* into these four groups (**Figure 3**). In comparison, these groups were less well supported by trees built from a single marker gene (full-length 16S rRNA), seven genes (MLSA) (10), or 120 genes (bac120)(36, 54). The same was true of the SNP-based phylogeny of the all *Streptococcus* genome set (**Supplementary Figures 2–5**). The accessory genome diversity closely mirrored taxonomic classifications (i.e. *S. mitis* group, *S. massiliensis*, *S. intermedius*, *S. constellatus*, and the taxonomic divisions within the SAC) (**Supplementary Figure 6**).

**Figure 2.**
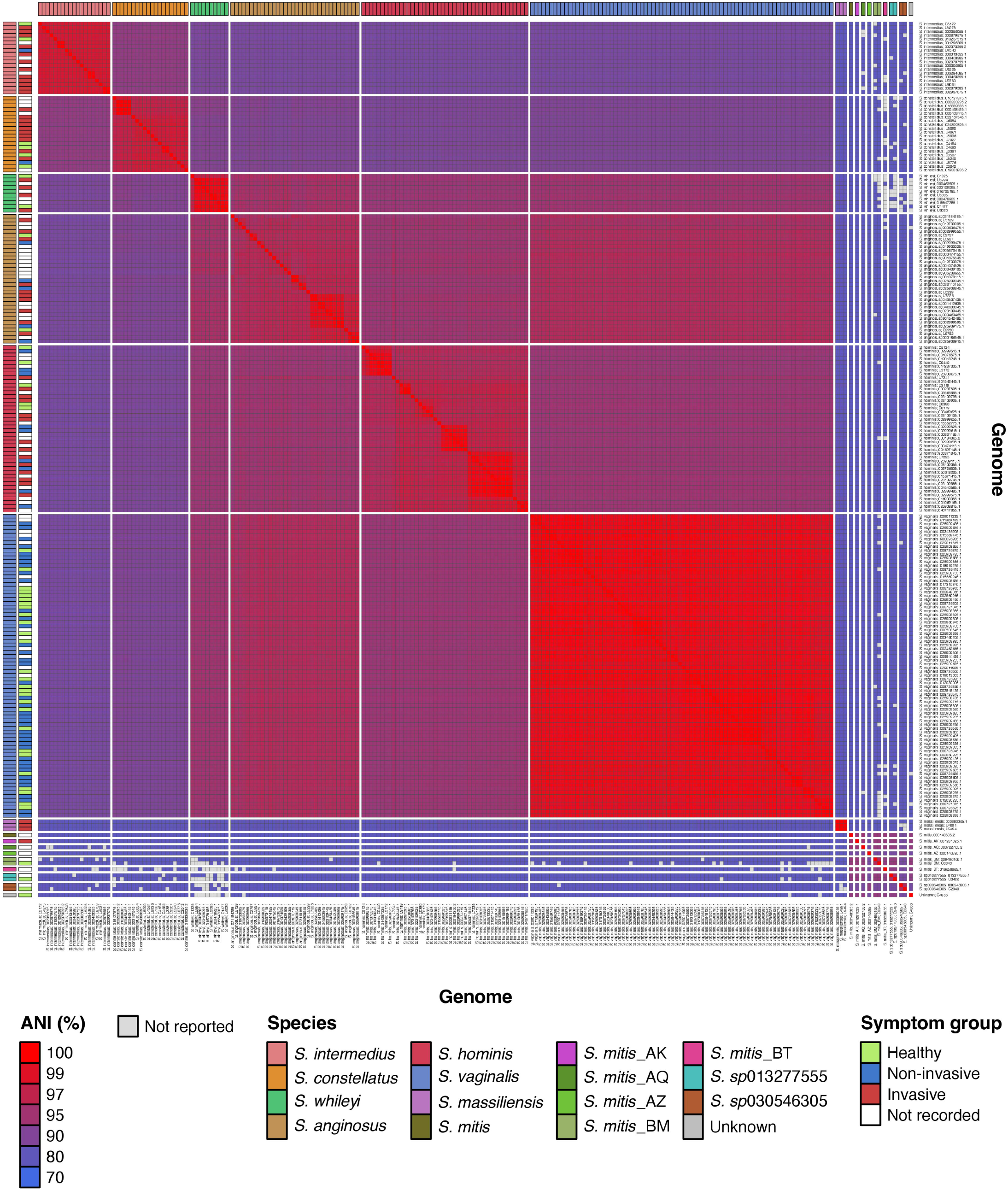
Average nucleotide identity across the de-replicated *Streptococcus* genomes. Pairwise average nucleotide identity (ANI) between all 222 de-replicated genomes (44 study isolates and 178 reference genomes). Each cell is the mean of the two fastANI directions, and grey cells fall below the fastANI reporting limit. Rows and columns are grouped by species, with the six SAG lineages ordered by invasive propensity (most invasive first) and then the other *Streptococcus* species. Strips show species and symptom group.

**Figure 3.**
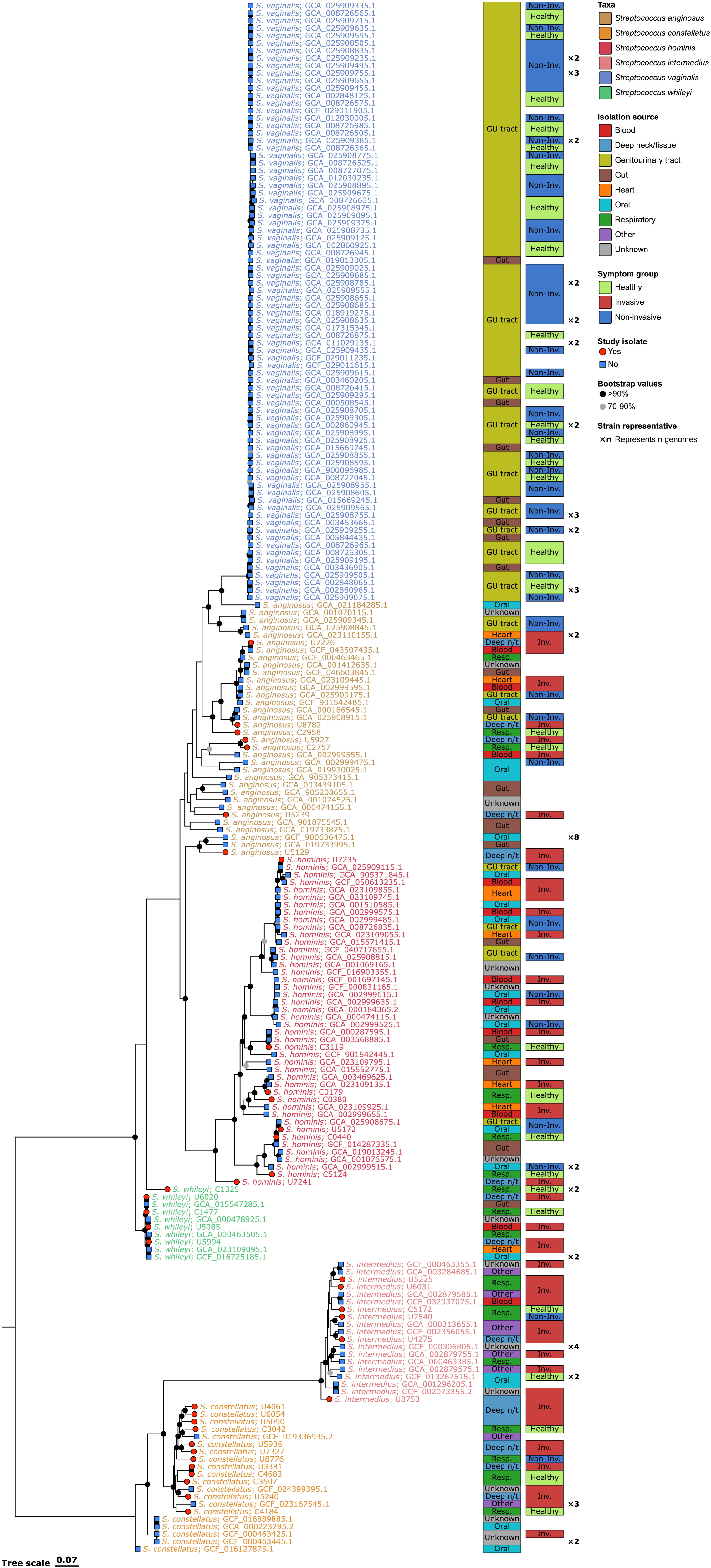
Core-genome SNP phylogeny of the *S. anginosus* group. Maximum-likelihood phylogeny (RAxML) of the 207 *S. anginosus* group (SAG) genomes, one per strain, inferred from SNPs in the Panaroo core-gene alignment. Rooted on the *S. intermedius*/*S. constellatus* clade. Leaf labels give the species and isolate or assembly accession, coloured by species; the two block columns show isolation source and symptom group (Healthy, Non-invasive or Invasive); red circles mark study isolates and blue squares reference genomes. Circles on branches show bootstrap support (black >90%, grey 70-90%). ×n marks a strain representative that stands for n genomes (strains: ≥99.9% average nucleotide identity and ≤10 cgMLST allele differences). Block abbreviations: Deep n/t = deep neck/tissue; GU tract = genitourinary tract; Inv. = invasive; Non-Inv. = non-invasive; Resp. = respiratory.

A discordance was noted in 10 genomes (three study isolates and seven reference) between their GTDB-Tk classification as *S. vaginalis* and their topology within the core genome SNP phylogeny tree, which placed them within the *S. anginosus* clade. Further investigation showed that the majority (6/10) originated from body sites *S. vaginalis* was not documented to be present in, as well as having all seven reference genomes being previously classified by Prasad et al. (17) as “Group 1” (i.e. the group which does not contain the *S. vaginalis* genomes) (**Supplementary Table 4**). Based on these observations, the genomes were reclassified as *S. anginosus* for the purposes of this study. Subsequent analyses with fastGEAR confirmed this classification decision by showing the genomes having a greater share of genes and ancestral sequences with *S. anginosus* rather than *S. vaginalis* (**Supplementary Table 4**).

Five participants (four hospitalised) samples yielded two isolates each. In three cases, MALDI-TOF and WGS produced discordant identifications (**Supplementary Table 2**). Within each of these three pairs, ANI exceeded 99.9%, indicating that the isolates were the same strain. Thus, one isolate from each pair was excluded from further analyses. Isolates identified as *S. massiliensis* or the *S. mitis* group were also excluded from comparative genomic analyses, as they fell outside the SAG focus of the study.

Because genome provenance metadata were inconsistently available, the remaining isolate and reference genomes were dereplicated to reduce the likelihood of analysing the same strain more than once. This yielded 207 SAG genomes (38 isolate, 169 reference) representing 206 strains (**Supplementary Tables 1** and **2**). Notably, eight of the 42 *S. anginosus* genomes were assemblies of the type strain deposited under six different strain designations. One strain comprising two *S. vaginalis* genomes from patients with different clinical outcomes was retained in full. Overall, cgMLST split 13 of the 31 SAG clusters defined by ANI alone.

#### 3.2.3 *S. anginosus* complex characterisation

To further resolve SAC membership and diversity, we evaluated ANI boundary separation and the equivalent separation based on patristic distances among the SAC genomes (n=168). These measures were interpreted alongside the topology from RAxML core-SNP trees at the all *Streptococcus*, SAG, and SAC levels, and with a SAC-level vertical inheritance tree inferred using Verticall (63) to account for recombination (**Supplementary Table 5**). Overall, SAC taxa formed distinct phylogenetic clades across the different taxonomic levels, including in the recombination-corrected tree (**Supplementary Figures 7** and **8**). The exceptions were *S. anginosus*, which showed unresolved, paraphyletic-like clustering across phylogenies, and *S. whileyi* which failed to form a monophyletic clade at the SAG level. Only the ANI boundaries of *S. vaginalis* and *S. whileyi* separated these taxa from their nearest neighbours (ANI gaps of +2.227 and +0.788, respectively), whereas both *S. hominis* and *S. anginosus* overlapped with theirs. Patristic distances showed the same pattern (**Supplementary Table 5**).

#### 3.2.4 Taxon cohesiveness is a gradient

Lineage cohesion was assessed using recombination-aware fastGEAR analyses of 1,251 gene alignments (**Supplementary Table 6**) from a SAC genome set dereplicated at 99.9% ANI (n=141). This gene panel represented approximately 60% of the SAC genome and was weighted towards core genes. Cohesion was defined as the mean proportion of shared sequence ancestry between genomes within a taxon (i.e. inherited similarity). *S. vaginalis* formed a highly cohesive lineage (98.7%) and *S. anginosus* the least (56.9%) (**Supplementary Table 7**). All four SAC taxa had cohesion values exceeding their respective permutation-based null expectations (n=9,999), indicating the positive coherence in each group occurred more than would be expected by chance. Across the 1,251 genes, *S. anginosus* shared nearly as much pairwise sequence ancestry with *S. vaginalis* (41.7%) and *S. whileyi* (44.4%) as within itself (56.9%) (**Supplementary Table 7**).

#### 3.2.5 The gene flow within the *S. anginosus* complex shows uneven donor:recipient dynamics

*S. vaginalis* and *S. hominis* were the taxa most involved in ancestral recombination within the SAC (33.1% and 31.1% of 2,074 ancestral events, respectively), and were also the highest donors of recent recombination events (**Figure 4**). However, *S. vaginalis* had by far the lowest recent ingression rate (2.3%) of any SAC member, resulting in it being the primary recent recombinant exporter (donor:recipient ratio of 15.5). The primary recipients from *S. vaginalis* were *S. anginosus* and *S. hominis*; nearest-neighbour comparisons indicated that the donated segments were more recent acquisitions, with median similarities to *S. vaginalis* of 99.72% in *S. anginosus* and 100% in *S. hominis*, compared with 91.35% and 90.55%, respectively, in unrecombined alleles. The abundance of *S. vaginalis*-derived segments did not differ between recipient genomes isolated from the genitourinary tract or gut (niche limit of *S. vaginalis*) and genomes from other sites (stratified permutation p = 0.82, 0.65 and 0.90 for genitourinary, gut, and both, respectively). This suggests that the segments were most likely acquired by ancestors of the sampled *S. anginosus* and *S. hominis* isolates that interacted with *S. vaginalis* in the genitourinary tract or gut. In contrast, *S. anginosus*, the SAC member isolated from the widest range of body sites, had the highest attributable inter-taxon recent recombination rate (53.4%), and was a net sequence importer (donor:recipient ratio 0.43).

**Figure 4.**
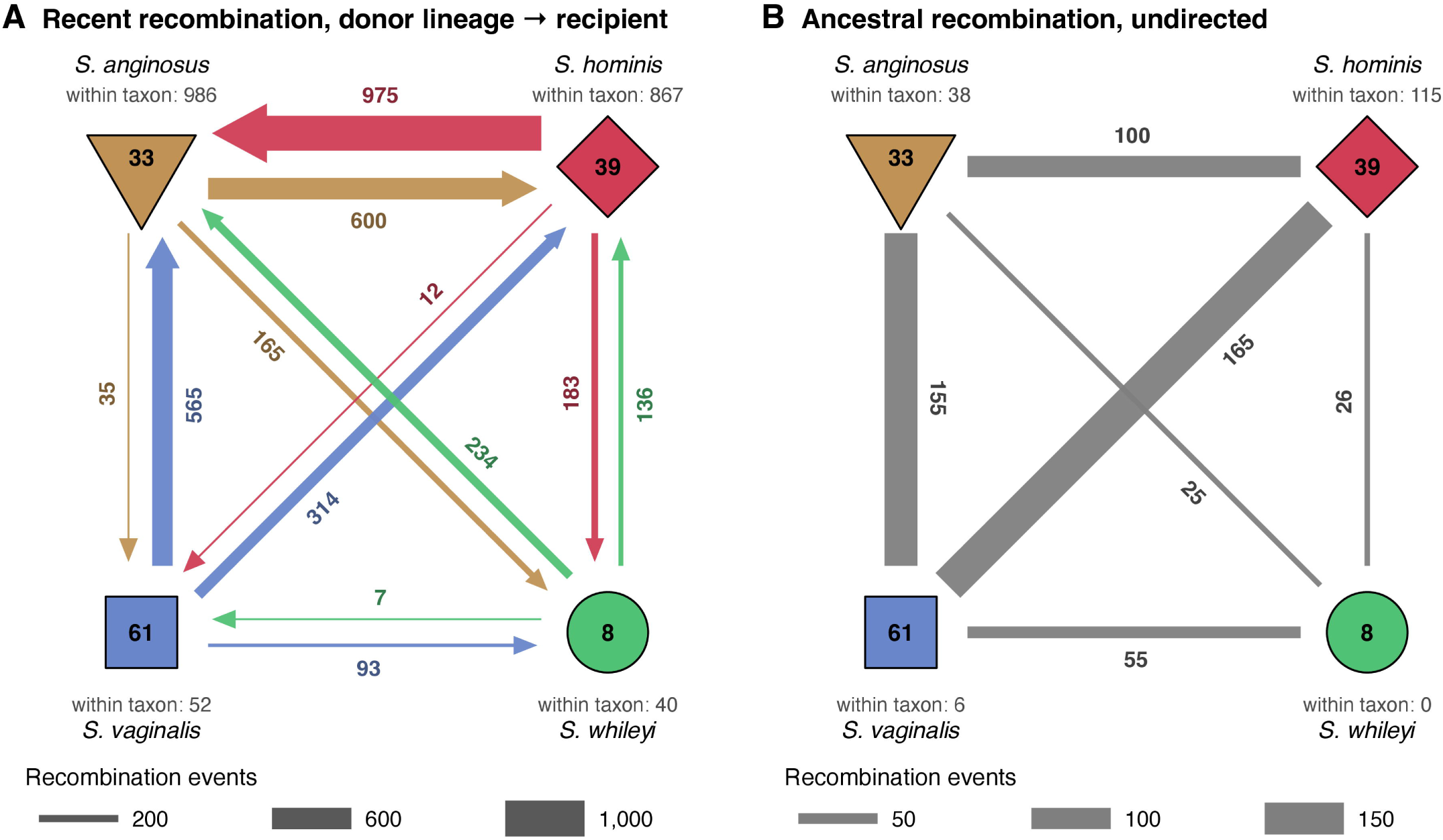
Recombination between the *S. anginosus* complex taxa has occurred unevenly over time. fastGEAR recombination across 1,251 genes in 141 clone-corrected genomes of the four *S. anginosus* complex taxa show *S. vaginalis’* heavy ancestral recombination, but minimised recent importation, contrasting with *S. anginosus* being the main importer of recent recombination. Nodes are taxa and show the number of genomes, with within-taxon events beneath. (A) Recent events from donor lineage to recipient taxon, with arrow width proportional to the number of events and arrow colour showing the donor taxon. (B) Ancestral events between lineages, drawn without direction, with line width proportional to the number of events (different scale from A). Recent and ancestral events are those after and before the most recent common ancestor of the affected lineage, and a recent event is counted once per recipient taxon.

### 3.3 Clinical profiles differ between taxa, especially within the *S. anginosus* complex

The SAG species distributions were similar between participants with invasive and non-invasive infections (**Supplementary Figure 9**, **Supplementary Table 2**) apart from *S. hominis* and *S. intermedius*. Five patients had polymicrobial infections; 80% were classified as invasive, which was similar to the proportion of those with monomicrobial infections (88.2%, 15/17) (**Table 2**, **Supplementary Table 2**). When including the reference genomes with available metadata, the propensity for invasive disease followed an overlapping gradient with *S. intermedius* showing the highest proportion of genomes associated with the invasive phenotype (80%) (**Figure 5**). The SAC members showed varying levels of invasive scoring, with *S. vaginalis* being the sole taxon which did not contain any cases of the invasive phenotype. Of note, *S. vaginalis* was also the only SAG taxon with a highly restricted body site range (**Figure 5**).

**Figure 5.**
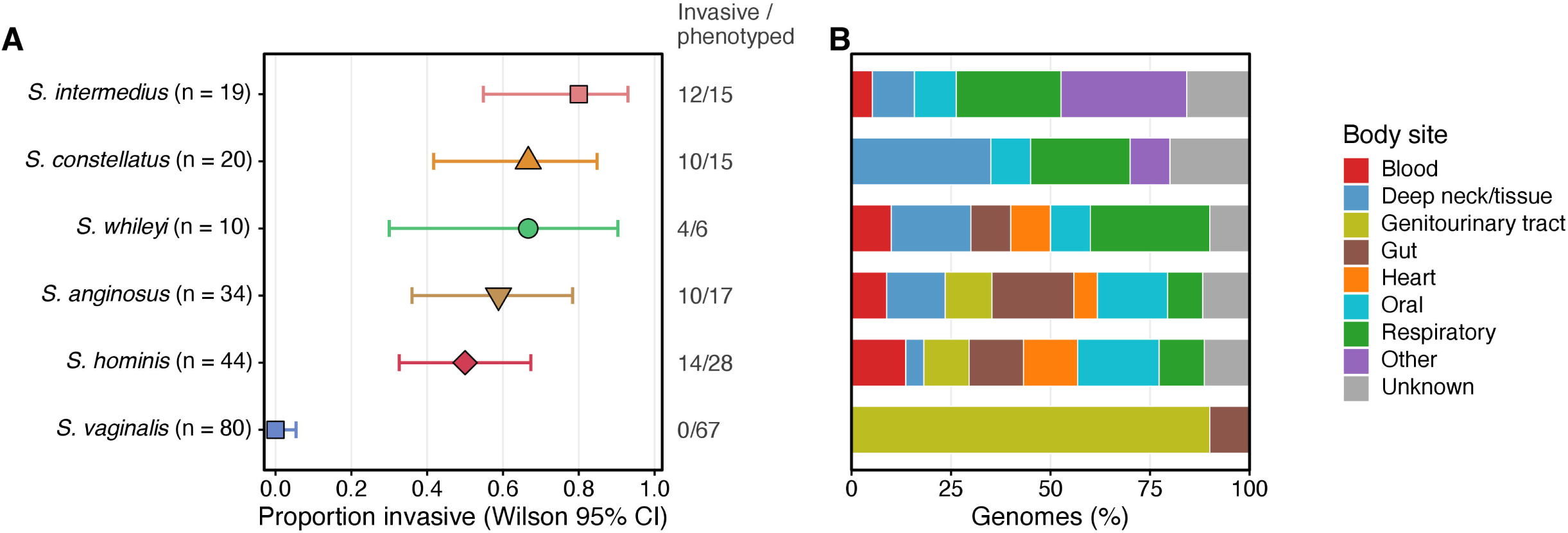
Invasive propensity and body site of isolation of the six SAG taxa. Increasing invasive propensity gradient and body sample site diversity within the combined isolate and reference genome cohort. (A) Proportion of phenotyped genomes (Healthy, Non-invasive or Invasive; n = 148) that are Invasive for each taxon. (B) Body site of isolation as the percentage of each taxon’s genomes (n = 207 de-replicated SAG genomes). Taxa are ordered by invasive propensity, most invasive at the top.

### 3.4 Functional characterisation

#### 3.4.1 The mobilome is primarily lineage-aligned

Prophage elements, all belonging to the *Caudoviricetes* class, were detected in 131 (63.3%) of SAG genome assemblies, with *S. vaginalis* exhibiting significantly higher prophage prevalence (78.8%; BH-adjusted p = 0.002) and *S. whileyi* the highest raw prevalence (90.0%, not significant after correction), whereas *S. anginosus* showed significantly lower prevalence (41.2%; adjusted p = 0.017). Prophage density differed between species (p = 0.027), with the highest means in *S. whileyi* and *S. vaginalis* (1.40 and 1.18 per genome, respectively, compared with 0.61–0.90 in the other SAG taxa), although no pairwise contrast remained significant after correction (**Supplementary Figure 10**). Prophage elements grouped into 81 viral clusters, most of which were highly specific; 56/81 (69.1%) were present only in single genomes and the majority (88.9%) being species-restricted (**Supplementary Figure 10**).

Overall, 83/207 (40.1%) of genome assemblies carried at least one plasmid element, with *S. vaginalis* having the highest prevalence (77.5%). Similar to the prophages, distinct plasmid clusters were primarily restricted to individual genome assemblies and were taxon-specific **(Supplementary Figure 10)**. Notably, no plasmids were detected in any of the study isolates. Further investigation of the individual assemblers’ contig outputs showed geNomad plasmid detections in assemblies from six isolates (**Supplementary Table 8),** all of which were subsequently identified as genome fragments containing conjugative elements, some of which spanned species and primarily carried restriction-modification, abortive infection and toxin-antitoxin cargo.

ISEScan detected 20 families of insertion sequences (IS) present across the SAG taxa (**Supplementary Figure 11**), with the lowest overall densities in *S. intermedius* and *S. anginosus*, and highest in *S. vaginalis and S. whileyi* (3.82, 8.41, 16.66 and 20.12 median copies/1000 CDS, respectively). Proximity of IS elements to function features varied, but broadly was more pronounced adjacent to DNA replication, repair and transcriptional regulation genes, and depressed in metabolism, transport and stress response genes (**Supplementary Figure 12**). Due to small sample sizes, taxon-specific enrichment category intervals were broad and could not be confidently assessed, apart from *S. vaginalis,* although this is likely a reflection of its larger sample size.

#### 3.4.2 Defence:CRISPR-Cas and prophage content are inversely related in all SAG taxa

DefenseFinder detected 70 distinct anti-phage defence system types across the SAG genomes, with a median of seven systems per genome (IQR 6–9). Total defence loads differed between species (per 1000 CDS, Kruskal–Wallis H = 64.7, p = 1.3 × 10 ^12^), with *S. constellatus* having the highest (median 5.06) and both *S. vaginalis* and *S. intermedius* the lowest (medians of 2.92 and 3.17, respectively) (**Figure 6**). CRISPR operons were found in 85 (41.1%) genomes, with most (n=54) carrying one operon, followed by 30 genomes carrying two, and a single genome carrying three operons. Multiple CRISPR-Cas subtypes could be found in most taxa, however no operons were detected in any *S. vaginalis* or *S. whileyi* genomes (**Figure 7**, **Supplementary Figure 13**). Fitting with expected biological mechanisms, within-taxon CRISPR-Cas and prophage were significantly correlated to each other in an inverse fashion for both prevalence and per-genome loads (**Figure 7**).

**Figure 6.**
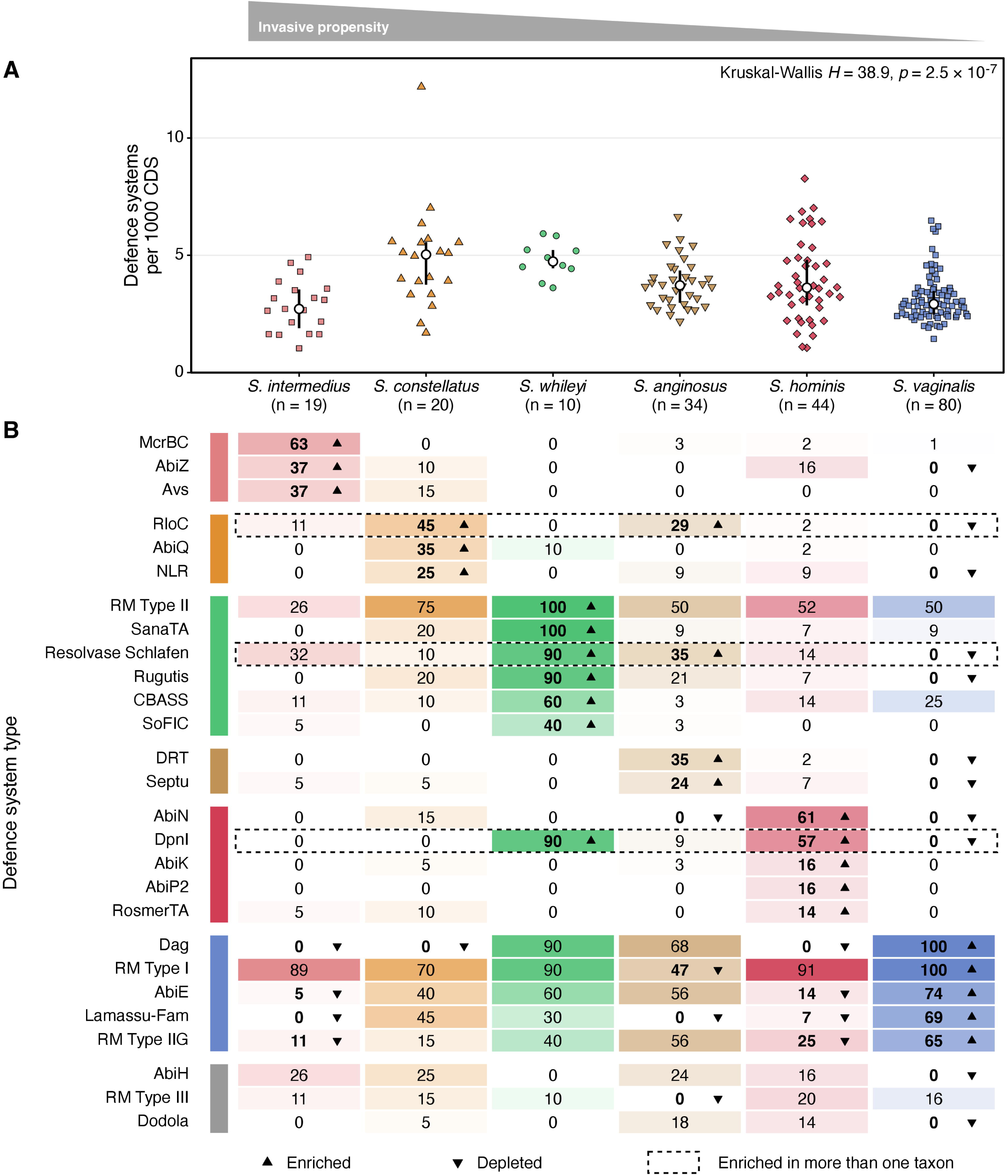
Non-CRISPR-Cas defence systems differ in load and composition between taxa. (A) Total non-CRISPR-Cas defence systems (DefenseFinder) per 1000 CDS per genome. Each point is a genome; white dot = median, vertical line = IQR (n = 207) (B) Defence-system types significantly enriched (upward triangle) or depleted (downward triangle) in a taxon versus all other taxa (Fisher’s exact test, Benjamini-Hochberg adjusted p < 0.05 over 414 tests); significant cells are in bold. Cells give the % of genomes carrying at least one system of the type, shaded from white (0%) to the taxon colour (100%). Rows are grouped by the taxon of strongest enrichment (coloured bar; 24 types) or as depletion-only (grey bar; 3 types), and a dashed outline marks types enriched in more than one taxon.

**Figure 7.**
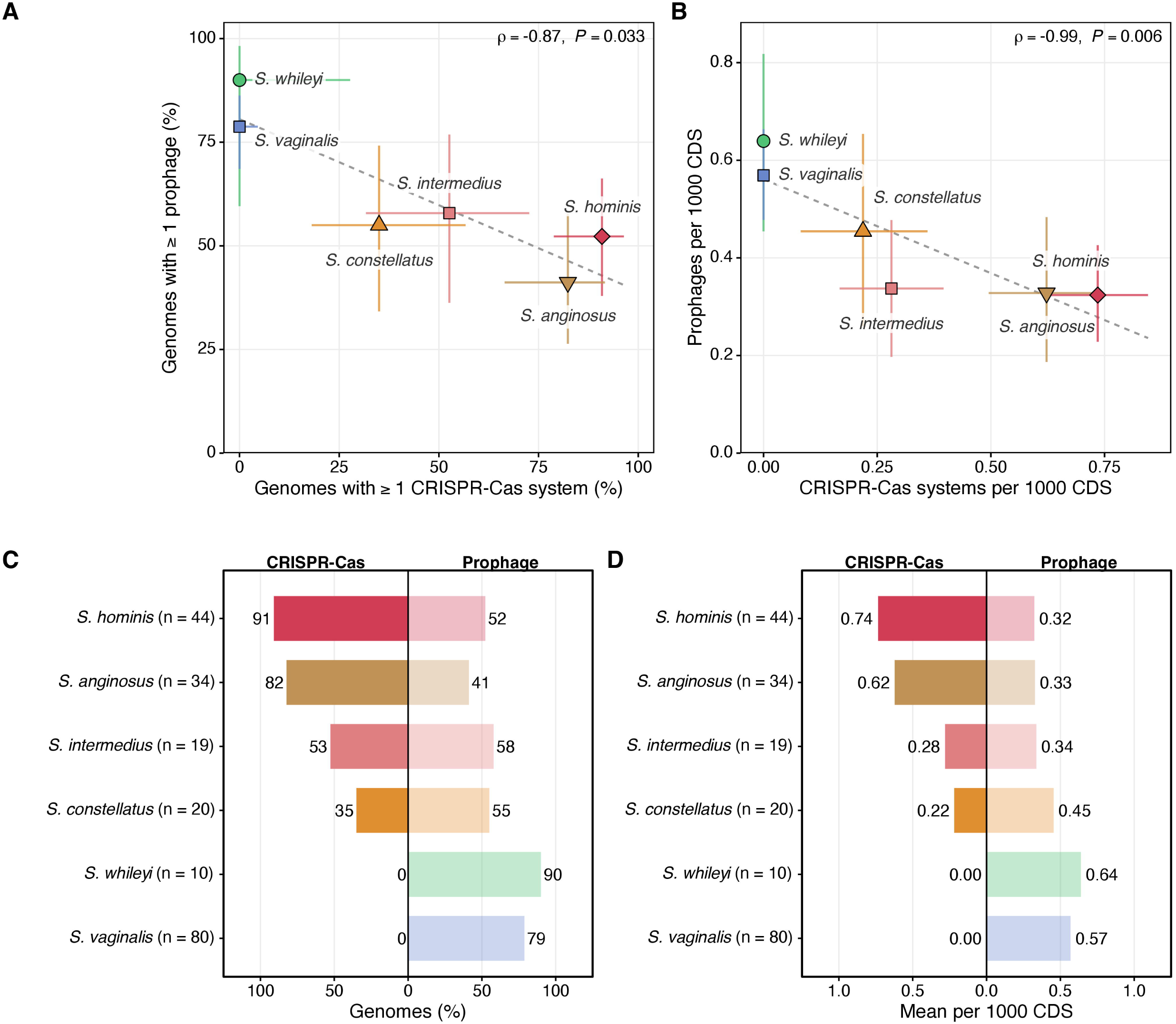
CRISPR-Cas content is inversely related to prophage content across the six SAG taxa. (A) Percentage of genomes with at least one CRISPR-Cas system against the percentage with at least one prophage, per taxon (n = 207 genomes); the point is the taxon percentage and the horizontal and vertical error bars are Wilson score 95% confidence intervals for the CRISPR-Cas and prophage percentages. (B) Mean CRISPR-Cas systems against mean prophages per 1000 CDS, per taxon; the point is the taxon mean and the error bars are 95% percentile bootstrap confidence intervals of each mean (10,000 resamples of the taxon’s genomes). (C, D) The values of A and B as back-to-back bars, with lineages ordered by CRISPR-Cas prevalence; CRISPR-Cas in solid colour, prophage in the lighter tint. Spearman rho with exact permutation P over the six lineages: A, rho = −0.87, P = 0.033; B, rho = −0.99, P = 0.006. Prophages are geNomad predictions of CheckV High-quality or Complete quality; the dashed line is a least-squares guide.

#### 3.4.3 Antimicrobial resistance profiles are taxa-linked but primarily driven by *S. vaginalis*

At least one AMR gene was detected in 14/30 (47%) of hospital cohort isolates compared to 2/8 (25%) of control cohort isolates, but after within-species permutation adjustment, the difference was not significant (p=0.46). With the inclusion of reference genomes, prevalence rates of genomes with ≥1 AMR gene ranged from 29% (*S. anginosus*) to 91% (*S. vaginalis*) (**Supplementary Table 9**). AMR gene density (genes per genome), resistance class counts per genome, and resistance class composition profiles differed significantly between taxa (**Figure 8**), with *S. vaginalis* contributing strongly to these differences (**Supplementary Table 9**). *S. vaginalis* was also the sole taxon containing Trimethoprim resistance elements (**Figure 8)**. To account for the larger sample size of *S. vaginalis*, the taxon was randomly downsampled to the median SAG taxon sample size (n=27), and the analyses were repeated over 200 draws. After downsampling, AMR gene density continued to differ significantly between taxa (median p<10^-7^), as did class and gene profiles (median R^2^ 0.137; p=0.002 and 0.138; p=0.002 respectively) (**Supplementary Table 9)**. For every AMR class detected in at least two genomes, *S. vaginalis* had the highest prevalence of any taxon, even after downsampling (**Supplementary Table 9)**.

**Figure 8.**
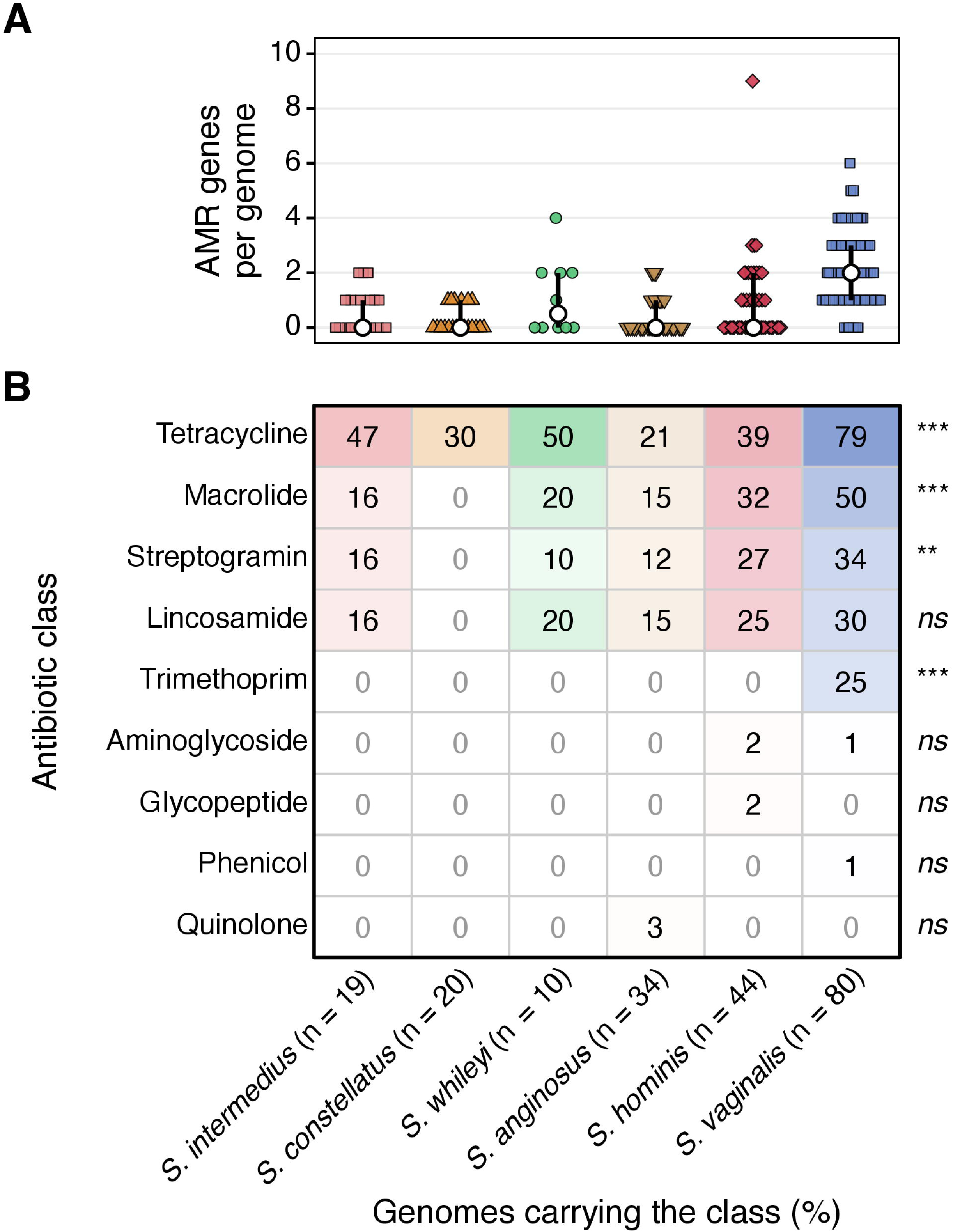
Antimicrobial resistance genes across the SAG lineages do not show resistance to standard antibiotic treatment. No beta-lactam resistance genes were detected, with overall resistance gene and class presence being most frequent in *S. vaginalis*. (A) AMRFinderPlus AMR genes per genome, with each point representing one of the 207 de-replicated SAG genomes. The white dot is the lineage median and the vertical line its interquartile range (no line where Q1 = Q3) (Kruskal-Wallis p = 4.2 × 10^-14; p = 0.12 excluding *S. vaginalis*). (B) Percentage of genomes carrying at least one AMR gene of each resistance class. The marks to the right of each row give Fisher’s exact test across lineages, adjusted for the 9 classes (* adjusted p < 0.05, ** < 0.01, *** < 0.001, ns not significant). n is the number of genomes per lineage; cells are shaded from white (0%) to the lineage colour (100%).

#### 3.4.4 Carbohydrate utilization profiles are lineage-oriented

Mean CAZyme family richness per genome ranged from 56.4 (*S. intermedius*), to 49.2 in *S. whileyi*, 47.6 in *S. constellatus,* 46.0 in *S. anginosus*, 45.0 in *S. hominis,* and 44.7 in *S. vaginalis*. Of these families, taxon differences were concentrated in 22 taxon-defining families within six modules (**Figure 9)**; all remaining families were variably shared across taxa. Ten families constituting a host N-glycan degradation cassette were present in all 19 *S. intermedius* genomes. Six of these (GH33 sialidase (NanA), CBM40, GH85, GH125, CBM16 and CBM47) were only found in *S. intermedius.* The remainder were variably found in *S. whileyi*, *S. constellatus,* and in a minority (2/80) of *S. vaginalis*. A second module comprising hyaluronate/chondroitin lyase (PL8), heparan sulfate lyase (PL12), unsaturated glucuronyl hydrolase (GH88) and the hyaluronan-binding module (CBM70) was exclusively present in all *S. intermedius* and *S. constellatus* genomes and in 9/10 *S. whileyi*. In contrast, a dietary galactan module was dominated by the presence (90–100%) of *S. anginosus*, *S. hominis,* and *S. vaginalis*, which was also the primary taxon for the starch and glycogen binding modules CBM20 and CBM34 (**Figure 9**).

**Figure 9.**
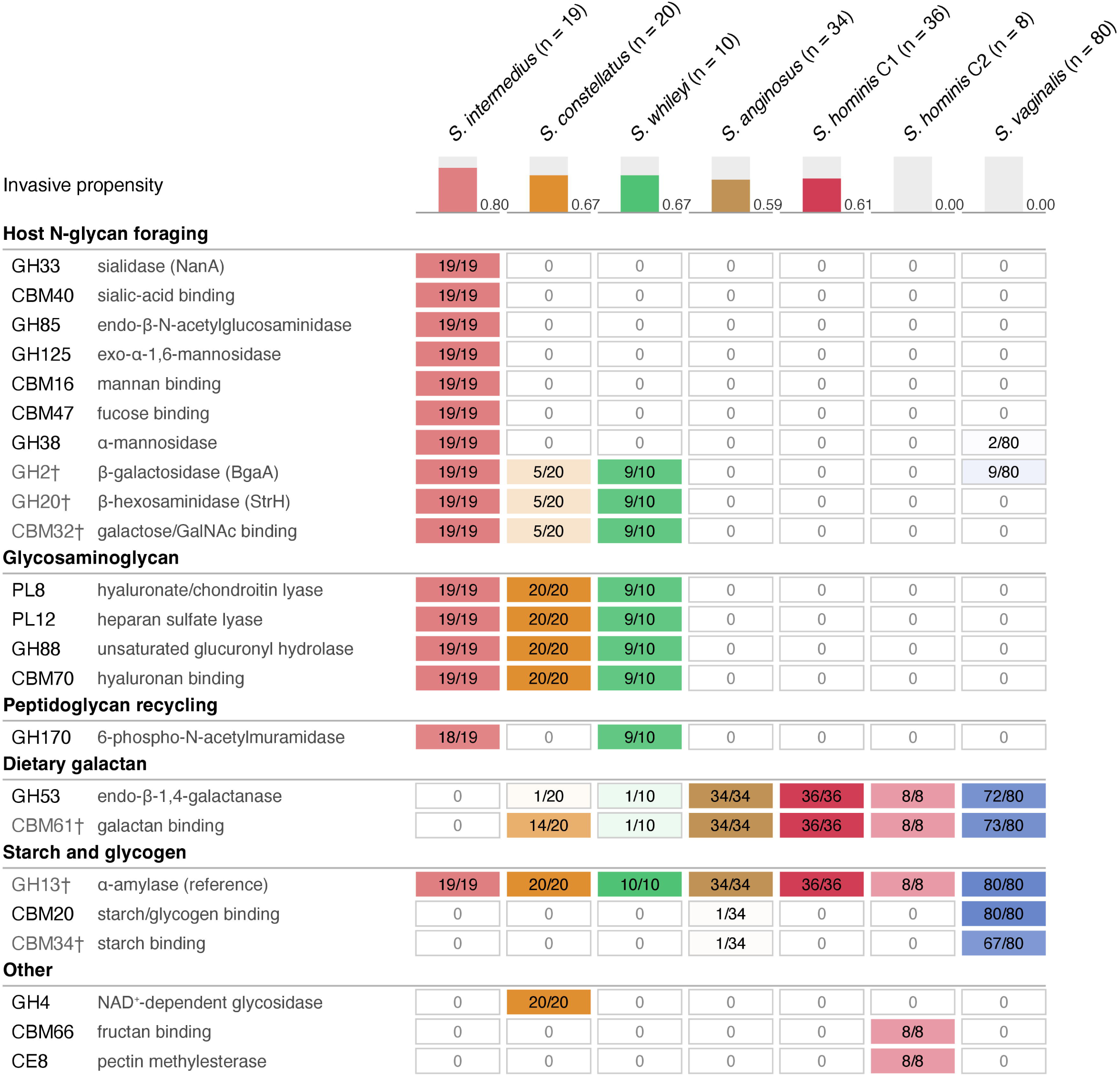
CAZyme families that separate the SAG taxa. Share of genomes per taxon carrying at least one gene in each dbCAN CAZyme family within the dereplicated SAG cohort (n=207). *S. hominis* is shown as its two clades, C1 and C2. Families are shown when at least one column carries them in 90% or more of its genomes and another in 10% or fewer, with families marked by a dagger being the exception and shown for context within their module. Families are grouped by predicted substrate module, and roles are homology-based.

#### 3.4.5 Sugar transporter systems are mostly conserved between SAG taxa

Bakta annotated 21–45 phosphoenolpyruvate:sugar phosphotransferase system (PTS) genes per genome, with many systems being universal (cellobiose, fructose, mannose) or near-universal (glucose, sucrose) between taxa. The mannitol PTS was only present in *S. vaginalis* (75/80), which also showed an expanded gene set for the galacticol PTS (3.7 genes/genome) compared to other SAG taxa (approximately 1 gene/genome). *S. constellatus* was enriched for the galactose PTS (70% vs 0 in other taxa) and in the β-glucoside PTS (95% vs ≤20% in other taxa). The N-acetylgalactosamine (GalNAc) PTS was present in all *S. intermedius, S. constellatus,* and in all but one *S. whileyi* genomes (**Figure 10**), which also perfectly aligned with the glycosaminoglycan (GAG) CAZyme module (**Figure 9**, **Figure 10**). Further analyses of the genomic regions encoding both features confirmed that they are adjacent to each other and form a GAG genetic cluster (**Supplementary Figure 14**), suggesting that the GalNAc PTS is the uptake step of the GAG degradation pathway. A single *S. whileyi* isolate (C1325) lacked the CAZyme families and PTS elements otherwise conserved in the taxon, and was the only isolate in the taxon with evidence of plant glycan degradation. Of note, C1325 was the representative genome of the 2-genome cluster falling outside of the main *S. whileyi* clade resulting in it failing its monophyletic clade test at the SAG level.

**Figure 10.**
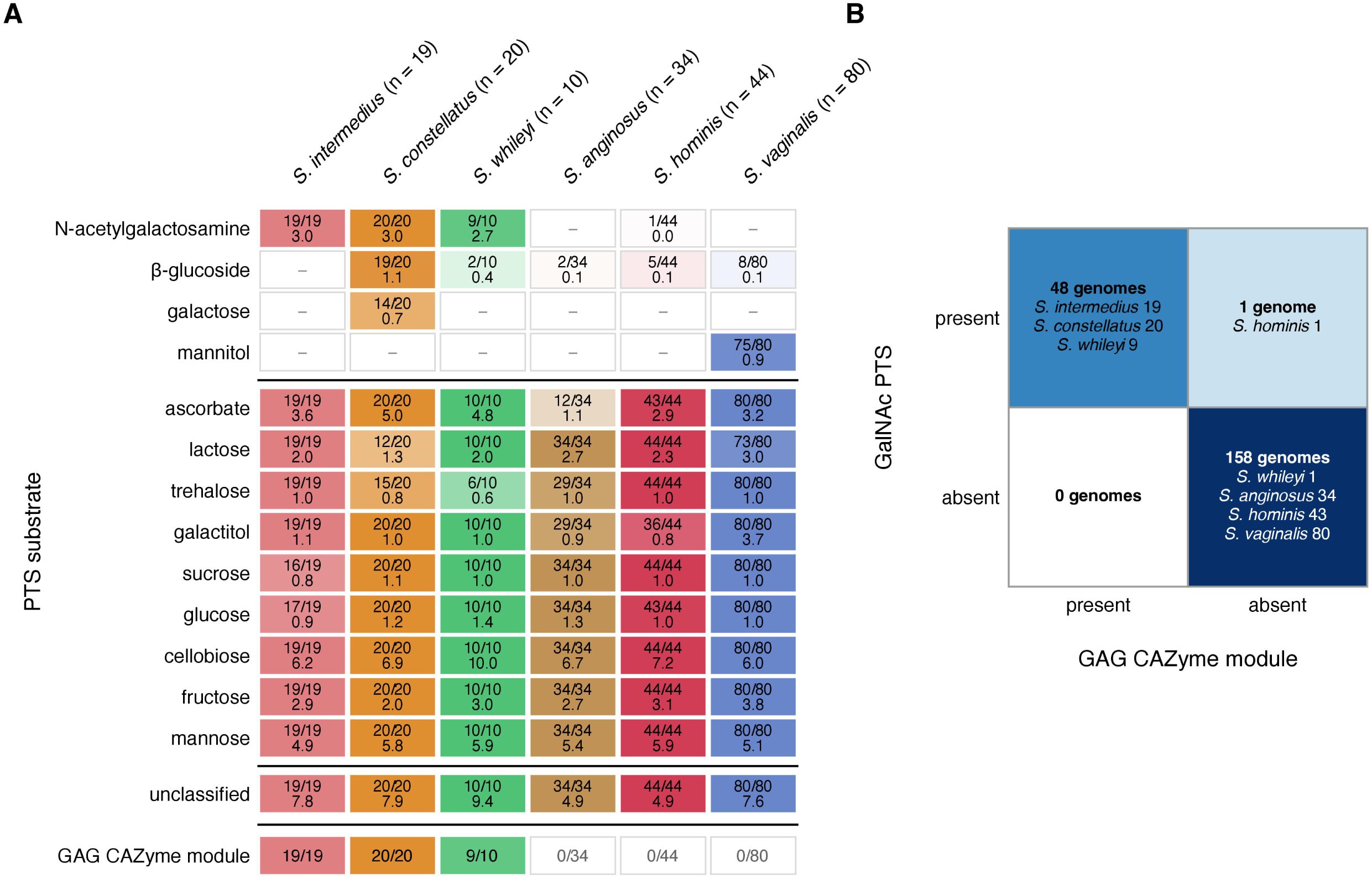
PTS substrate predictions and the GalNAc PTS x GAG CAZyme module. (A) PTS substrate carriage, with cell values showing the number of genomes carrying at least one gene for the substrate over the total taxon genomes on the top, and mean genes per genome at the bottom. Unclassified = PTS genes with no substrate match, dash = no genes detected. Columns ordered by taxon invasive propensity (n=207). The bottom row gives the number of genomes carrying the GAG CAZyme module (PL8, PL12, GH88 and CBM70 all present). Lineages are ordered by invasive propensity (n = 207 genomes). (B) Co-occurrence of the GalNAc PTS and the GAG CAZyme module showing a taxa-associated pattern of the two linked elements: 48 genomes carry both, 158 neither, and one in which only a GalNAc-named PTS gene was detected.

#### 3.4.6 Virulence factor profiles are sparse and act as lineage markers

Virulence factor (VF) load was low and largely fixed by taxon, ranging from a mean of 8.0 VFDB hits per 1,000 CDS in *S. intermedius* to 4.7 in *S. vaginalis* (**Figure 11, Supplementary Table 10**). Of note, VFDB *Streptococcus* representation does not include any of the SAG species, so database matches represent homologues and naming conventions may not translate precisely. The VF composition varied by taxon, with consistent differences including *S. intermedius* exclusively carrying *nanA* neuraminidase (sialidase/GH33) and lacking *rfb*A LPS (monoderms such as *Streptococcus* do not have LPS; likely *rml*A which helps supply rhamnose for cell wall polysaccharide synthesis), both *S. whileyi* and *S. vaginalis* lacking capsule genes, and the low number of VF hits overall in *S. vaginalis*. Intermedilysin, a hallmark VF of *S. intermedius*, was not listed because VFDB lacks SAG representation, however its *ily* gene was detected in the Panaroo annotation of all *S. intermedius* isolates and no other taxon.

**Figure 11.**
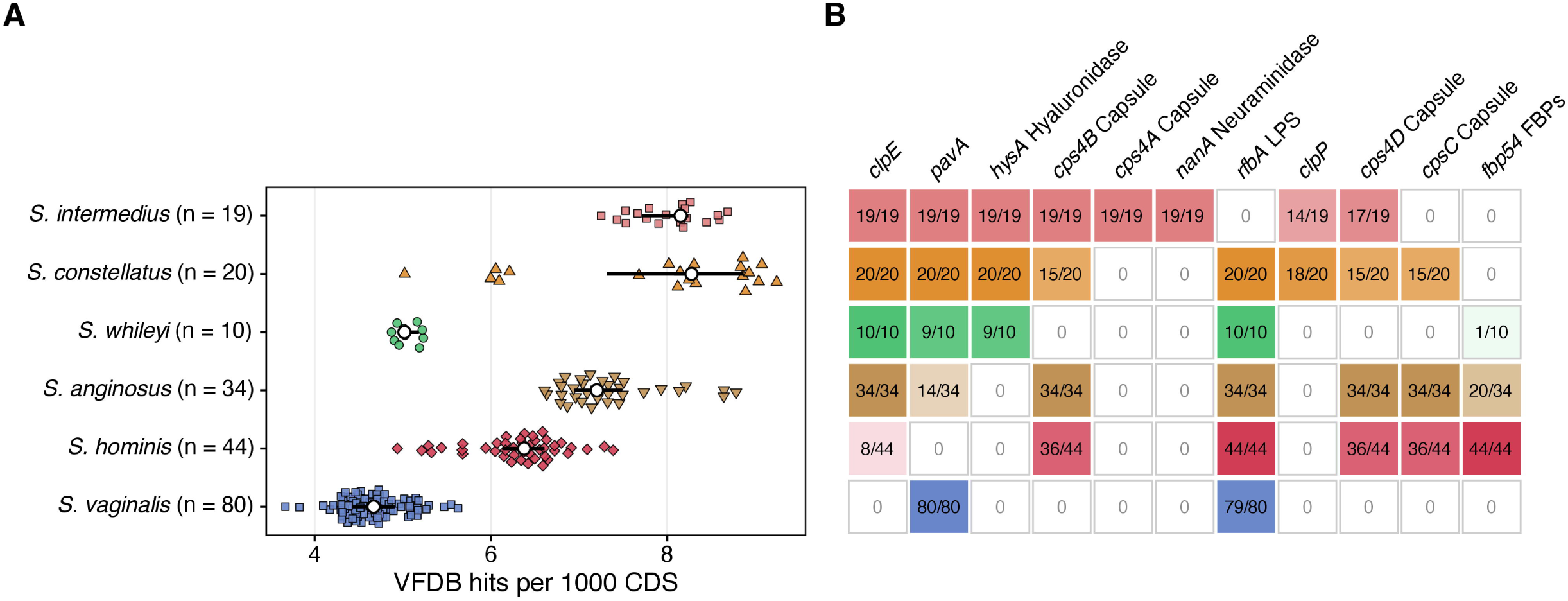
Virulence Factor gene loads and compositional differences between SAG taxa. Virulence-factor genes (VFDB) in the 207 de-replicated SAG genomes. (A) VFDB hits per 1000 coding sequences, with each point representing one genome. /=The white dot is the median and the line the interquartile range (Kruskal-Wallis p = 4.3e-35). (B) VFDB genes carried by at least 90% of genomes in one taxon and at most 10% in another (11 of 29 genes detected). Rows are aligned to taxa in panel A, and cells give carriers over total taxon genomes, shaded from white (none) to the taxon colour (all). Taxa are ordered by invasive propensity, most invasive at the top.

#### 3.4.7 A small number of metabolic modules differentiate the SAG taxa

Genome-scale metabolic reconstruction was performed with gapseq on 206 of the 207 genomes, with one exclusion due to frameshift errors within the tested genomic blocks (**Supplementary Table 1**). The 583 variable gene-backed reactions and transporter substrate calls were collapsed by exact presence-pattern identity into 222 blocks (**Supplementary Table 11)**. Per-genome block richness formed two tiers: 94.7 blocks in *S. constellatus* and 97.8 in *S. intermedius* against 108.9 to 110.4 in the other four lineages (**Figure 12**). Most (124/165) of the reconstructed transporter substrate calls were shared, with the small number of differentiating block patterns being taxon-specific (**Figure 12**). Of note, a 36-feature butyrate fermentation block was present in every *S. anginosus*, *S. constellatus*, *S. whileyi* and *S. hominis* clade 1 genomes, but absent in *S. intermedius*, *S. vaginalis* and *S. hominis* clade 2 genomes (see 3.4.8). Mannitol and α-galactosidase usage was near-exclusive to *S. vaginalis,* as was its near-absence of siderophore biosynthesis which contrasted with the siderophore’s ubiquity across all other taxa (**Figure 12**).

**Figure 12.**
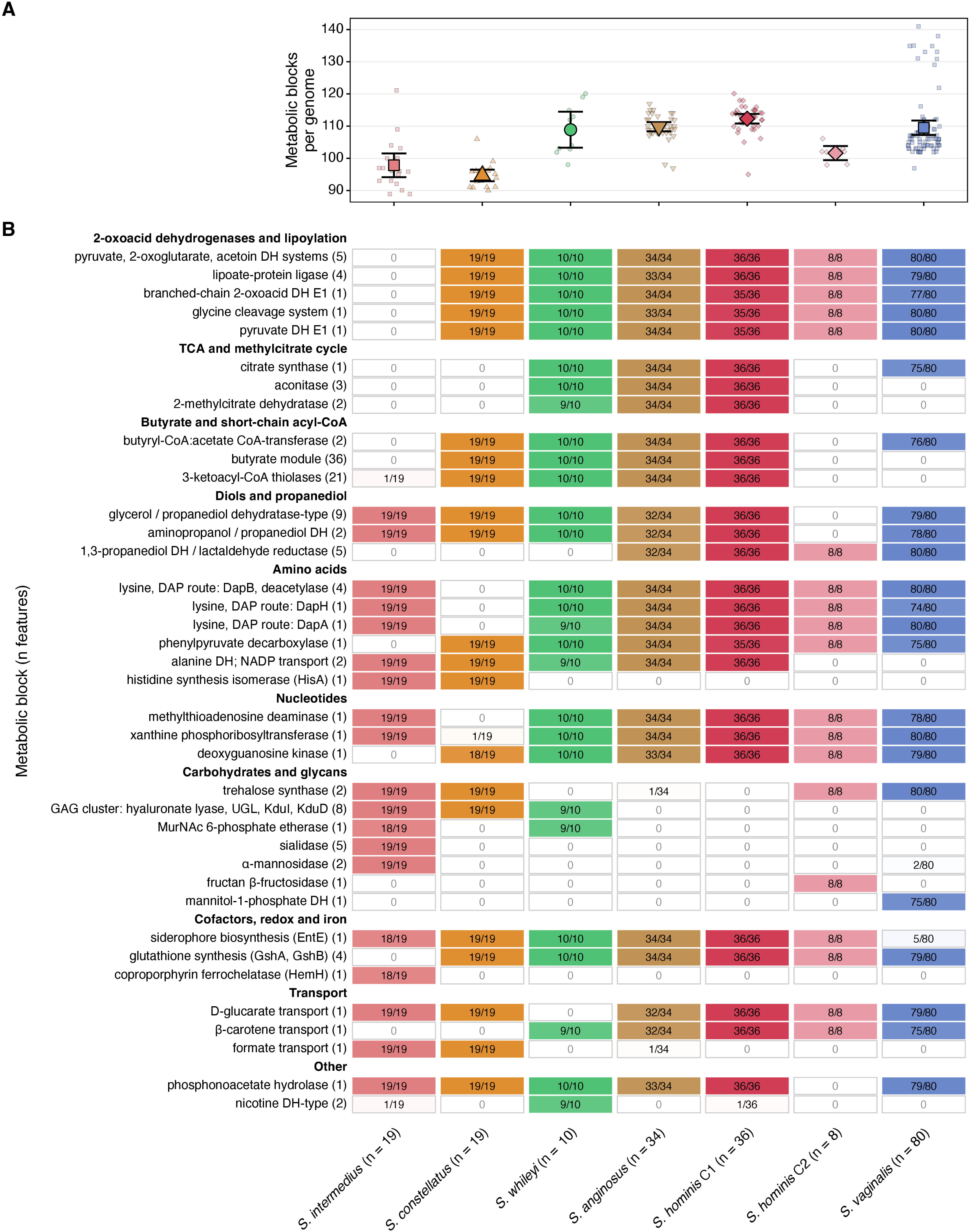
Metabolic block counts and composition differ by SAG taxa. Metabolic blocks (sets of gapseq features with identical presence across genomes) in 206 de-replicated SAG genomes. (A) Blocks carried per genome. Large symbols are lineage means with 95% confidence intervals and small symbols are individual genomes. (B) The 38 blocks for which every lineage or clade carries the block in at least 90% or at most 10% of its genomes. Cells show carriers / total genomes in the taxon, shaded from white (none) to the lineage colour (all), and the number of features in each block is in brackets. *S. hominis* is shown as its two clades, C1 and C2. The *S. constellatus* denominator is 19 because GCA_000223295.2 has no gapseq reconstruction.

#### 3.4.8 *S. hominis* has two phylogenetically and functionally distinct clades

The *S. hominis* taxon consistently resolved into two clades in both the core SNP and Verticall phylogenetic trees (**Figure 3**, **Supplementary Figures 2–5**), prompting further investigation. The larger Clade 1 (n=36) contained the *S. hominis* type strain (GCA_014287335.1) but appeared paraphyletic, while Clade 2 (n=8) formed a monophyletic offshoot. Mean ANI between the clades was 96.13%, above the commonly used species-level. Despite this, the clades still differed across unrelated functional systems: the butyrate fermentation block, aconitase and the *cps* capsule locus were present in all Clade 1 and no Clade 2 genomes, whereas the *eps* capsule locus (8/8 vs 20/36), α-galactosidase (5/8 vs 0/36), an additional fructan-active protein made of a GH32 catalytic domain flanked by two CBM66 binding modules (**Figures 9**, **11**, **12**) and the AbiP2 anti-phage defence system (7/8 vs 0/36; Fisher’s exact p = 2.1 × 10) were enriched in Clade 2.

### 3.5 Invasive correlations

#### 3.5.1 Accessory gene and core gene GWAS does not detect correlation with invasive phenotype

Genetic associations with invasiveness were tested in pyseer using accessory gene presence/absence and core-genome SNPs. *S. vaginalis*, which contributed no invasive genomes, was excluded. In the SAG genomes (66 genomes: 50 invasive and 16 non-invasive), no accessory gene presence/absence pattern was significantly associated with invasiveness. The lowest p-value was 1.2 × 10 ³, approximately 14-fold above the Bonferroni significance threshold, with little evidence of genomic inflation (λ = 0.90). No core-genome SNP reached significance either, despite the more stringent Bonferroni threshold reflecting the larger number of unique SNP patterns tested. Using the kinship matrix from a recombination-corrected phylogeny (Verticall) in place of the core-SNP phylogeny left the gene-level result unchanged (**Supplementary Figure 15**).

#### 3.5.2 Invasive potential is a property of lineage and not locus

After accounting for lineage, none of the tested genomic features was significantly associated with invasiveness, including nucleotide identity, core-gene SNP distance, insertion-sequence composition, functional axis content, accessory gene presence/absence, and the 222-block metabolic space. Insertion-sequence family composition was the only feature to retain nominal significance after lineage adjustment (p = 0.038), but this association reached p < 0.05 in only 6.1% of *S. vaginalis* down-samples, suggesting that it was sensitive to cohort composition (**Figure 13)**. At the gene presence/absence level, however, the three taxa with the highest invasive propensity all contained glycosaminoglycan-degrading loci and GalNAc-annotated phosphotransferase systems, with one exception among the genomes. These features co-occurred with the GH88 gene in a single approximately 15 kb cluster within the PTS operon. This cluster was absent from the other three taxa, which had lower invasive propensity or were non-invasive (**Figure 13**). A 36-feature butyrate fermentation block was present in all SAG taxa except *S. intermedius* and *S. vaginalis*, which represented the extremes of invasive propensity. Its absence alone cannot be considered a determinant of invasive potential. Intermedilysin, sialidase, and a host N-glycan cassette were present in all of the *S. intermedius* genomes, however these features were taxon-fixed (i.e. act as markers of lineage rather than invasiveness potential) (**Figure 13**). Across the six taxa, Spearman correlations between invasive propensity and seven feature categories (virulence factors, defence systems, insertion-sequence families, prophage clusters, CAZyme families, PTS substrates, and AMR gene richness) identified nominal associations with AMR gene richness (ρ = −0.928, exact p = 0.017) and insertion-sequence family richness (ρ = −0.899, p = 0.028). However, neither association met the Bonferroni-adjusted significance threshold of p < 0.00714 (**Figure 13**).

**Figure 13.**
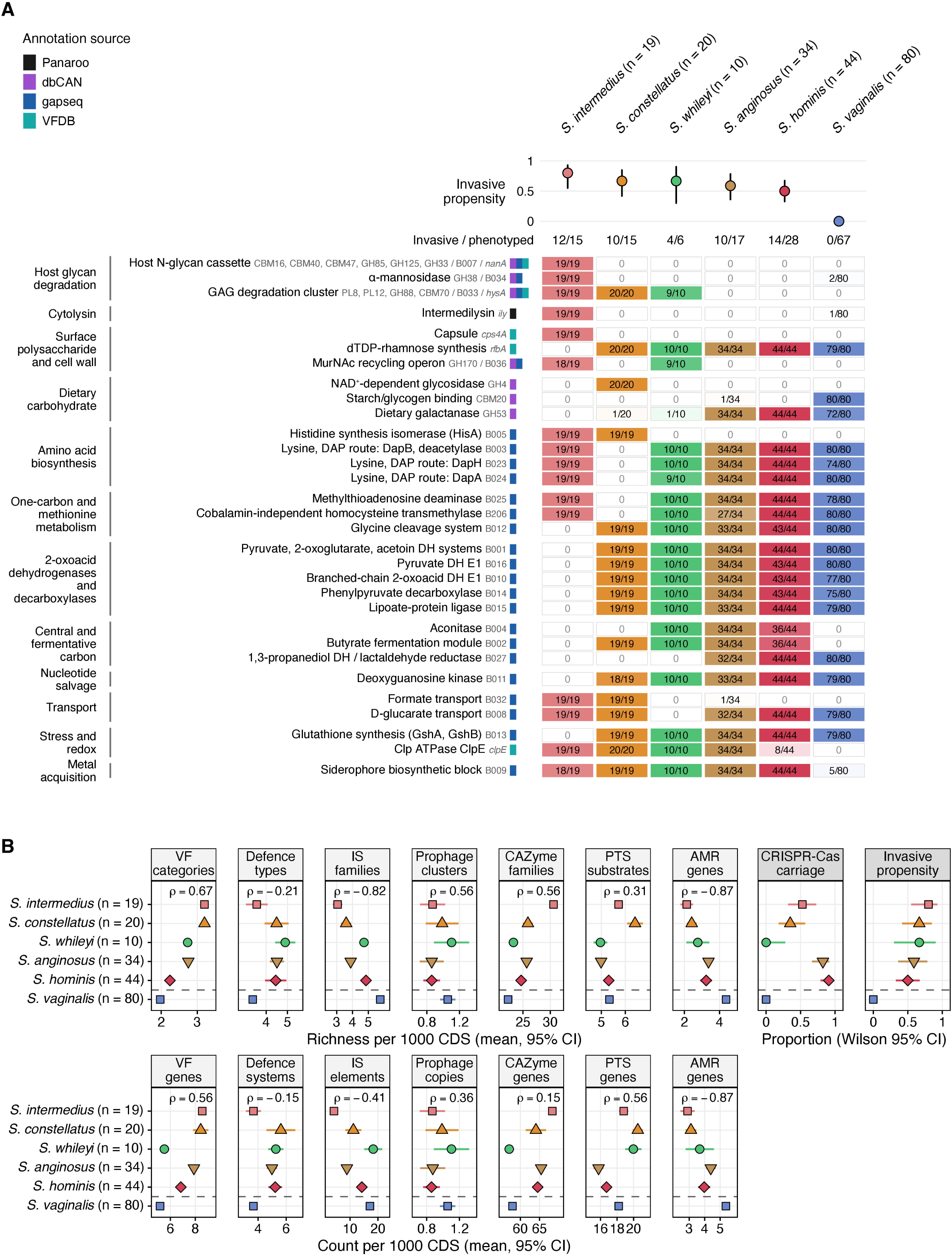
Taxon-discriminating genomic features and functional richness of the SAG taxa against invasive propensity. Host-glycan degradation was the only functional theme whose distribution followed the severity ordering rather than being defined by lineage. (A) Functions carried by at least 90% of the genomes of one taxon and at most 10% of another, as determined by Panaroo gene presence/absence, dbCAN CAZyme families, gapseq metabolic blocks and VFDB genes (all n=207 apart from gapseq n = 206). Annotations from different tools that mark the same function, with identical presence in every genome, are merged into one row; the features are listed in grey after the row name (gapseq blocks as B numbers of the Figure 12 block set) and the coloured squares give the tools. Rows are grouped by functional theme; cells show carrying / total genomes in taxon; invasive propensity (proportion of phenotyped genomes that are invasive, Wilson 95% interval) is shown above each taxon with the invasive / phenotyped counts. (B) Per-taxon means with 95% confidence intervals of seven genome-content features per 1000 coding sequences (top row, richness; bottom row, abundance). Spearman rho against the proportion invasive is computed over the five taxa above the dashed line (*S. vaginalis* has no invasive phenotyped genome), with an exact permutation p; CRISPR-Cas carriage and invasive propensity are proportions with Wilson intervals and are not tested. Taxa are ordered by invasive propensity.

## 4 Discussion

The original aim of the study was to characterise local SAG isolates to enhance diagnostic capabilities. However, unexpected discrepancies between routine species identification methods and the WGS data prompted a broader investigation into the diversity of SAG members, their predicted functional capabilities, evolutionary and population dynamics, as well as potential impacts on clinical outcomes. Furthermore, streptococci have complex genetic histories, including broad capacity for horizontal gene acquisition (82, 83). Genomic analysis of the SAG can therefore clarify how its member taxa have adapted to the human host and what roles they play. This study illustrates that the conventional concept of the SAG is more complex than currently recognised and highlights the value of WGS for accurate species identification and improved characterisation of SAG diversity.

### 4.1 MALDI mis-identification rate is high compared to genomic sequencing

In many clinical laboratories SAG are detected using culture-based methods supplemented with MALD-TOF species identification. Aside from the limitations of culture-based methods (84), our data demonstrated poor correlation between identification on MALDI-TOF and WGS classification, with agreement in only 42.8% of isolates. This applied not only to closely related species within the SAG group, but also included *S. massiliensis, S. mitis* group (including a *Streptococcus* uncharacterised as the species level) and *Pauljensenia odontolytica.* While MALDI-TOF diagnostic specificity issues are known, generally the rate of disagreement is lower than what is reported in this study (85), and likely due to our use of the more specific WGS-based identification rather than PCR/MLSA, as well as the subdivision of the *S. anginosus* species into four taxa classified as “species” by GTDB-Tk. Indeed, if the GTDB classifications of the SAC taxa were to be collapsed into a singular *S. anginosus* group, then the concordance rate would increase, albeit only to 69.4% (n=34).

In this study, we found that comparative genomics analyses benefited from high-quality, single-contig genome assemblies through long-read sequencing, which capture a more accurate and less fragmented genome architecture than what conventional short-read sequencing typically delivers (86). Previous work has highlighted the importance of single-contig genomes for accurate SAG comparative genomics (87). Beyond assembly quality, phylogenetic resolution improved with increasing numbers of loci, progressing from a single locus (16S rRNA), to seven housekeeping genes (MLSA), to 120 marker genes (GTDB), to core-genome SNPs. However, for finer resolution of groups falling mostly inside the 95% ANI similarity window, the threshold conventionally used to define a species, phylogenetic analyses benefitted from focusing on specific *Streptococcus* groups (e.g. the SAG). This is partly due to the increase in number of core genes able to be sampled for generation of the discriminatory SNP loci. Using this layered classification strategy allowed us to reveal a complex and diverse genomic narrative within the SAG.

### 4.2 SAC taxa classification

We found that the genomic diversity of the conventionally classified *S. anginosus* was much greater than that of *S. intermedius* and *S. constellatus*, in line with previous reports (10, 17, 87). This diversity has been recognised in proposed sub-species (subsp. *anginosus*, *whileyi* and genomosubsp. AJ1) (10), and, more recently, in a “Group 2/Urinae” (17). However, the lack of clear separation between these groups at the conventional >95% ANI species threshold (55) makes it harder to support splitting them into separate species. The GTDB classification scheme also assigns species using ANI (36). This makes an otherwise robust method prone to errors where species have indistinct or overlapping ANI boundaries. Such overlap can explain why GTDB-based classification could not distinguish some *S. anginosus* genomes that lay in the boundary overlap with *S. vaginalis*, even though, on tropism, gene content and phylogenetic topology, these genomes were closer to *S. anginosus* than to *S. vaginalis*.

Based on the study analyses, we propose that the conventional *S. anginosus* species is a “complex” of four closely related but distinct taxa which align with previously described *S. anginosus* subspecies; *anginosus* (10), *whileyi* (10), the recently proposed “Group 2 / Unrinae” (16, 17) and a clade which corresponds to the type strain of *S. hominis* (88). GTDB named the Group 2 / Urinae genome group “*S. vaginalis*” as it clustered with the previously described type strain of *S. vaginalis* (81); consequently this nomenclature was chosen for use in this study as well.

*S. vaginalis* was the best defined lineage of the complex, with the highest within-lineage nucleotide identity, the shortest within-lineage patristic distances relative to its nearest neighbour, the highest fastGEAR cohesion of the four taxa (98.7% shared ancestry), a near-complete absence of an otherwise SAC-conserved siderophore biosynthesis block, total loss of the *cps* capsule locus, and a conserved, taxon-specific (i.e. ≥95% present within taxa, ≤2% present in other taxa) metabolic complement that includes an exclusive mannitol PTS operon and 22 accessory gene clusters. Both *S. vaginalis* and *S. hominis* formed monophyletic clades at all test levels, however unlike *S. vaginalis*, *S. homonis*’ boundary gaps were negative for both topology-free measures (−0.404 ANI, −0.026 patristic). Its within-taxon cohesion was also lower (65.8% shared ancestry), however this is primarily driven by its division into two distinct clades which are both structurally and functionally differentiated, alongside only Clade 1 containing invasive-associated genomes. The *S. whileyi* taxon had clearly defined, non-overlapping boundaries commensurate with its high taxon cohesion (88.3% shared ancestry), however it failed on one of 5 clade tests, which was due to the singular genome C1325 that contained a taxon-divergent functional profile (e.g. absence of otherwise conserved host N-glycan). Other functional features unique within the SAC membership were however fully conserved across the taxon (e.g. complete loss of both the *cps* and *eps* loci), or features only shared with *S. vaginalis* (e.g. total absence of a CRISPR-Cas system). *S. anginosus* as defined in this study had the broadest within-taxon ANI spread (3.295), alongside having an overlapping ANI boundary of 1.283 with *S. vaginalis*, compared to *S. hominis* which shared a much smaller (0.404) overlap with *S. anginosus*. The within-taxon diversity and ANI overlap were consistent with a lack of monophyletic clustering at any phylogenetic level and the absence of taxon-specific accessory genomic content. The *S. anginosus* within*-*taxon cohesion (56.9%) however did clear its permutation null (46.3%), and contained taxon-conserved features contrasting to those of *S. vaginalis* (e.g. presence of a butyrate module, *cps*, etc..) despite the two taxa’s closest proximity by topography and ANI to each other. Indeed, if examined across the feature space, *S. anginosus*’ holistic “footprint” appears much more similar to *S. hominis* (**Supplementary Figure 16**); consequently its relation to other SAC members appears contingent on what assessment framework is used: it is closer to *S. vaginalis* by ANI, closer to *S. hominis* by composite features, has the highest shared recombinant ancestry fraction with *S. whileyi* (44.4%).

### 4.3 Plasmids, IS elements and evolution

*S. vaginalis* has effectively withdrawn from recent recombinant exchange while participating in at least a third of attributable ancestral events, making the restriction recent, with a defence-mediated explanation (i.e. prevention of entry or degradation of exogenous DNA) unlikely given its minimal restriction-modification core and its high plasmid carriage. Thus, ecological isolation following restriction to the genitourinary tract is the more parsimonious scenario, matching the habitat-specific recombination described by Shapiro et al. (89). Direction and timing ratios support this: *S. vaginalis* donated 15.5 segments and 18.0 events for each one received. Crucially, those segments lay within the donor lineage’s own nearest-neighbour diversity and so are recent copies rather than retained ancestral variation, and most events reached two or more sampled genomes, implying ancestral acquisition on a shared mucosal surface followed by inheritance rather than ongoing transfer at the recipients’ isolation sites. The taxon’s lack of adaptive anti-phage (e.g. CRISPR-Cas) systems while retaining tuneable, sequence-free permissiveness (e.g. Dag, restriction modification families) to various mobile elements fits its top ranking for prophage carriage amongst the SAGs, and alongside its high plasmid and CDS-normalised IS element loads demonstrate that the taxon is not closed off to all genetic flow; just to the recombination mechanism. These lines of evidence, in combination with *S. vaginalis* also having the largest genome of the SAC taxa, support a mobilome-driven genome expansion and are the expected population-genetic signature of relaxed purifying selection within a reduced effective population size following niche restriction (i.e to the genitourinary tract) (90, 91). A high mobilome load however is not a general marker of recombination restriction and better tracks population history. For example, *S. whileyi* carries an equal or greater burden compared to *S. vaginalis* while remaining an active participant in recent recombination. At the other end of the spectrum is *S. anginosus*, the most recombinogenic taxon and a net recipient, with the lowest IS element and prophage and second lowest plasmid loads, and no conserved taxon-specific accessory content. These metrics track *S. anginosus*’ borderline lineage cohesion, indicating that either this is a fuzzy taxon maintained by continuing exchange across a porous boundary (92) or a nomenclatural container of undescribed lineages driven by insufficient sampling. Across the SAC, common cross-lineage homologous exchange alongside largely taxa-specific prophage pools indicates a shared recombination pool but separate mobile-element pools. Unfortunately, there were insufficient genomes with appropriate metadata to estimate a date or pattern of speciation using molecular clock techniques.

### 4.4 Comparative genomic and metabolic specialization across the *S. anginosus* complex

Streptococcal carbohydrate acquisition occurs in large part through phosphotransferase systems, which import and phosphorylate their substrate in one step, as well as linking carbohydrate availability to virulence gene expression (93, 94). Together with the secreted and surface glycosidases, the PTS repertoire help predict a lineage’s accessible nutrient range **Supplementary Figure 16**)..

*S. intermedius* encodes the complete enzymatic machinery for sequential degradation and utilization of host N-linked glycans, alongside a glycosaminoglycan-degradation locus, while lacking the capacity to degrade dietary plant galactan. This repertoire may favour exploitation of continually replenished epithelial glycans as a relatively stable nutrient source, while glycan-degrading enzymes may also remodel host glycoproteins and expose underlying adhesion receptors, potentially contributing to invasive risk(95–97). Commitment to one reliable substrate is matched by a coherent reduction elsewhere: *S. intermedius* alone lacks the 2-oxoacid dehydrogenase and one-carbon metabolic features fixed across the rest of the group, giving it the second-smallest reaction network, almost no mobile content, and the largest conserved taxon-specific accessory complement. This functional distinctiveness has thus been vertically maintained rather horizontally acquired.

*S. constellatus* contained the same host glycosaminoglycan-degrading module as *S. intermedius* on the narrowest reaction network in the set (94.7 blocks per genome) but lacked the sialidase-led N-glycan cassette. Its β-glucoside and galactose transport capability therefore sits downstream of glycosidases it does not itself encode, suggestive of glycan cross-feeding dependencies in polymicrobial oral or respiratory communities (98–100) rather than purely relying on its own independent catabolic capabilities. *S. constellatus* had the highest innate defence count per CDS paired with almost no CRISPR-Cas and no plasmids. This profile is consistent with an organism with a relatively stable genome and niche adaptation that is also surrounded by a diversity of competitive threats, such as the oral and respiratory tract environment.

*S. whileyi* sits in the middle of the group, rather than at the peripheries. It carries the host-glycan degradative module that otherwise marks the abscess-forming lineages *S. intermedius* and *S. constellatus*, with GH2, GH20, CBM32 and the MurNAc-recycling operon (GH170/murQ/MurP, 9/10) shared almost exclusively with *S. intermedius*. In contrast to the former two SAG taxa, it also has retained the large reaction network of the other SAC taxa (e.g. aconitase and butyrate fermentation blocks), the former at 10 of 10 alongside *S. anginosus* and *S. hominis* clade 1. That composite is matched by its ecology: six named isolation sources, against three to four for *S. constellatus* and *S. intermedius* and seven for *S. anginosus* and *S. hominis*, so the lineage recovers the body-site breadth of the generalists while retaining the degradative apparatus of the specialists. Thus, *S. whileyi* is functionally a lineage linking the restricted, small-network abscess formers to the broader-niche, large-network members of the SAC.

The substrate repertoire of *S. hominis* is directed at dietary rather than host polysaccharide: it carries none of the glycosaminoglycan-degrading module found in the abscess-forming lineages (PL8, PL12, GH88 and CBM70 at zero across all 44 genomes) and none of the N-glycan cassette, while retaining endo-β-1,4-galactanase and its galactan-binding module (GH53 and CBM61 at 100%) on one of the larger reaction networks in the complex. This combination of broad fermentative capacity on diet-derived substrates accompanies its distribution across the widest range of body sites recorded in the SAG taxa (seven named isolation sources jointly with *S. anginosus*), as would be expected of a lineage whose nutrient base is available at any colonised mucosal surface rather than at a particular tissue. Within the species that range is partitioned rather than uniformly occupied: the two clades differ reciprocally in fermentable substrate (the butyrate module and aconitase at 36/36 against 0/8, α-galactosidase at 0/36 against 5/8) and in surface polysaccharide system (cps 36/36 against 0/8, eps 20/36 against 8/8), which points to a nuanced generalist model of two populations of distinct nutritional and surface biology.

*S. anginosus* combines the commensal-end substrate profile with the most complete core fermentative repertoire in the complex: it is missing the glycosaminoglycan module and the N-glycan cassette, but retains dietary galactan degradation, the butyrate fermentation module and aconitase. Thus, the taxon’s functional range lies in core metabolism rather than in specialised degradative capacity directed at host tissue. It carries N-acetylneuraminate lyase at 33 of 34 genomes without any sialidase, pointing towards a dependence on other microbial community members to liberate the host sialic acid it consumes. Alongside *S. hominis*, its functional flexibility is matched by the joint-widest range of recorded body sites of isolation. The ability to maximize flexibility appears to come with a cost however, with both *S. anginosus* and *S. hominis* encoding the largest diversity and density of CRISPR-Cas systems, as well as high densities of defence system elements.

*S. vaginalis* is the only SAG taxon equipped to draw on glycogen, a host storage polysaccharide, with the distinction of the mechanism being a matter of capture rather than catalysis. The taxon’s α-amylase is the same GH13 glycol-degrading enzyme carried by every SAG genome in the collection, but it alone pairs that enzyme with *S. vaginalis*-fixed starch and glycogen binding modules (e.g. CBM20/34). The relevance of such an arrangement appears in context of its preferred urogenital environment being *Lactobacillus*-dominated: many *Lactobacilli* utilise glycogen-derived sugars, but cannot catabolise glycogen itself, and so are reliant on host α-amylase to extracellularly hydrolyse the glycogen(101). Thus, being able to capture and directly degrade the glycogen into usable catabolic derivatives gives *S. vaginalis* a competitive advantage in accessing an abundant carbon source, and mirrors a similar convergent strategy used by another co-resident of the environment, *Lactobacillus crispatus* (102). Around that core the taxon has traded the host-glycan and glycosaminoglycan machinery its relatives retain for a SAG-exclusive mannitol operon (75/80 genomes) which would allow it to exploit high local levels of mannitol found in association with *L. crispatus* (103), suggesting the presence of a cross-feeding mechanism.

These features, alongside expanded galactitol transport and near-fixed α-galactosidase, align with previous reports of “*S. anginosus*” isolates from genitourinary tract samples having species-atypical, expanded fermentation capabilities (e.g. mannitol and raffinose), such as the description by Bergman at al (104). Indeed, a blast search (16S_ribosomal_RNA database) of the provided short 16S rRNA gene sequence in the Bergman et al. article confirmed the presumed classification; a singular near-identical hit (98%) to the *S. vaginalis* type strain P1L01 (NR_179340).

Of note, the urogenital niche’s glycogen enrichment is thought to be driven by estrogen induced accumulation in the vaginal/cervical epithelium, which is then liberated into the luminal pool through exfoliation and lysis (105). Furthermore, the diet-based polyol degradation modules speak to the vaginal-perineal-distal gut anatomy continuum, consistent with 8/80 genomes originating from stool. In contrast, the male urogenital tract offers neither on a comparable scale: glycogenated epithelium is transient and confined to a distal anterior penile area(106), there is no standing mucosal fluid pool, and dietary-derived plant sugars do not reach the tract at all. The prediction is therefore that *S. vaginalis* has evolved to primarily reside in the female urogenital tract, and the *“S. anginosus*” that is detected in the male urethra is likely transfer from vaginal intercourse (107). Correspondingly, most, if not all *S. vaginalis* genomes with available metadata to date have been sourced from female subjects irrespective of study origin (16, 81).

### 4.5 Invasiveness

In our analysis there were no locus-level, accessory gene, or core-genome SNP associations with invasive infection. The lineage-adjusted analyses support this interpretation. Nucleotide identity, SNP distance, functional feature content, accessory gene presence and the 222-block metabolic space all lost significance after lineage adjustment. At the presence/absence level, the taxa most associated with invasive disease (*S. intermedius*, *S. constellatus*, *S. whileyi*) carried a ∼15 kb PTS-associated cluster combining putative glycosaminoglycan-degrading loci, GalNAc-annotated phosphotransferase systems and GH88. Streptococci are known to catabolize host glycosaminoglycans: extracellular lyases depolymerise hyaluronan and chondroitin sulfate, the resulting unsaturated disaccharides are imported through dedicated PTS transporters, and the GH88 unsaturated glucuronyl hydrolases release the unsaturated uronic acid, which is then converted to intermediates of central metabolism (108, 109). This capacity could contribute to invasive disease by providing a carbon source in host tissue, where free sugars are limited, and by facilitating tissue penetration through degradation of extracellular matrix. However, because the 15Kbp PTS cluster tracked taxa and lineage rather than invasiveness within lineages, we interpreted it as a taxon-level trait that may predispose to invasive potential, rather than as a determinant of invasive infection in our cohort. We found intermedilysin, sialidase and the host N-glycan cassette to be conserved within *S. intermedius*. Intermedilysin is a human-specific cytolysin and a well-characterised virulence factor of this species (110) but its taxon-wide fixation means it marks the lineage rather than explaining variation in invasive propensity. Similarly, the 36-feature butyrate fermentation block was absent from both extremes of the invasive spectrum (*S. intermedius* and *S. vaginalis*), so its absence cannot be invoked as a causal invasiveness factor. Likewise, *S. whileyi* is the only taxon in the group to have neither capsule system (*cps* 0/10, *eps* 0/10) at a severity propensity of 0.667, which also eliminates capsular polysaccharide presence as a lineage-independent risk of increased invasive potential.

Taken together, invasive potential as measured in this study was found to be an emergent property of functional profile composition rather than a property of any singular locus or feature, and these functional profiles typically aligned with lineage. Specialised profiles geared towards active host-substrate liberation, and thus taxa which had such profiles (e.g. *S. intermedius, constellatus,* and *S. whileyi*) had the highest invasive potential, while conversely, the SAG taxa which had the broadest functional and catabolic profiles tended to be less prone to invasion.

### 4.6 Antimicrobial Resistance carriage in SAG

The AMR carriage was higher in hospital isolates compared to non-hospitalised (47% vs 25%), although this was not significant after within-species permutation. AMR gene density, resistance class counts and class composition differed significantly between taxa, with *S. vaginalis* having the highest prevalence (91%) and being the only taxon with trimethoprim resistance elements, which may reflect more frequent exposure to antibiotics as they are excreted in the urine. The most common AMR genes detected in the study were against tetracycline and macrolides, however these finding are more of an evolutionary question of retention versus fitness cost rather than being of clinical relevance, as beta-lactams are typically the first line treatment of choice in SAG infections (5, 6). Differential profiles and rates of AMR gene carriage between the SAG taxa, and their different genetic diversity acquisition strategies, however, means the potential for evolving clinically-relevant AMR in one or more taxa remains a possibility.

### 4.7 Non-SAG infections presenting as SAG-associated disease

Although not the focus of this study, it is noteworthy to mention the detection of *S. massiliensis* in two patients hospitalised with invasive disease, which despite near-identical (ANI >99.9%) genomes, were identified by MALDI-TOF as *S. constellatus* and *S. anginosus*. This mis-identification is likely due to *S. massiliensis* not being represented in the MALDI-TOF database used for the original diagnosis. *S. massiliensis* has been rarely documented in the literature beyond the original published case of a single subject in 2006 isolated from blood in a patient with pneumonia following surgery for a gunshot wound (111). Yet in our investigation, we found the species in 9% (2/22) of cases with invasive disease, suggesting that *S. massiliensis* may be more common and clinically important in head and neck pathologies than previously recognised.

Additionally, a MALDI-TOF classified SAG isolate was identified as *Pauljensenia odontolytica* by GTDB. This species was reclassified from *Actinomyces odontolytica* and is termed by NCBI as *Schaalia odontolytica*. It has been reported as an early colonizer in multispecies oral plaque (112) and in one reported case of bacteremia (113). Our isolate was obtained from intracranial pus as part of a polymicrobial infection with *Eikenella corrodens* and *Capnocytophaga gingivalis* and may either be a sampling contaminant or genuine co-infection originating from the oral cavity.

### 4.8 Outlook and implications from the study results

Our findings have several practical implications for research and clinical practice. First, type strains cannot be assumed to represent their taxa; the predicted nutrient and metabolic variation we observed implies that assumptions built on a single reference strain may not be generalisable across the group, as exemplified by the *S. vaginalis* type strain (P1L01) being in the minority (5/80) of the taxa’s genomes to not contain the key mannitol fermentation module. Second, some caution should be employed in the utilisation of public genome databases. As was the case encountered in this study with the *S. anginosus* type-strain, multiple uniquely-named genome assemblies can in fact be duplicate copies of the same strain, which if not accounted for, will inflate apparent sample sizes and weaken the independence assumptions of downstream statistics (114). Misclassification is a larger problem. When GTDB-Tk was applied to all available GenBank genome assemblies annotated as *S. anginosus* which were not generated by this study (n=534) (**Supplementary Table 11**), the dominant taxon turned out to be *S. vaginalis* (61.6%, n=329), followed by *S. hominis* (20.2%, n=108), *S. anginosus* (15.5%, n=83) and *S. whileyi* (2.1%, n=11). Because *S. vaginalis* is site-restricted, carries more AMR genes and is less pathogenic, its dominance under the traditional “*S. anginosus*” label could be a major confounder in any analysis linking species, species features and clinical outcomes, and could distort diagnostic evaluations. For example, a traditional view of the “*S. anginosus*” species would pool multiple taxa whose invasive propensity ranged in this study from 0 to 0.67. Careful classification considerations are therefore advised prior to undertaking any such analysis. Finally, a minority of hospitalised cases were polymicrobial infections, however these data were derived from routine culture-based diagnostics, which have limited sensitivity for polymicrobial infections. Metagenomic sequencing is superior at detecting polymicrobial infections, with such infections being shown to be associated with more severe disease (115). Therefore, relying on conventional culture-based diagnostic methods may lead to an underestimation of the true rate of polymicrobial infections in SAG cases. This is relevant for both clinical management (e.g. triaging for higher risk of severe infections), and in the analysis of the mechanistic etiology of the invasive phenotype if it is either wholly or partly a microbial community property (i.e. cultured SAG genome-only association analyses will be blind to the important co-infection covariates).

### 4.9 Limitations

Our study had limitations including test sample sizes that were small in some situations. Depending on the context of the analysis, the ability to draw conclusions was impacted due to hard limits to available genomes for representative taxa, availability of metadata, or from potential sampling bias, and as such, the findings in this study should serve as guideposts for more in-depth research. We noted that no plasmids were detected in our sequenced SAG isolates, despite the generation of high-quality full-chromosomal assemblies. This may be a genuine outcome as our isolates belonged to no or low-plasmid prevalence SAG taxa, however we cannot exclude technical loss of small plasmids due to ligation-based ONT library effects (116). The gene patterns described in this study are associative in nature and need to be further investigated before any can be considered as demonstrated determinants of invasive capacity.

### 4.10 Conclusions

In conclusion, the SAG are much more complex than the three recognised taxa. Current clinical identification using MALD-TOF may be insufficient for accurate species-level identification, particularly if considering *S. hominis, S. vaginalis,* and *S. whileyi.* We found that WGS-based identification using GTDB in conjunction with core-gene phylogeny to be superior to MALDI-TOF, 16S or MLSA and should be considered the new gold standard in the research setting. The invasive potential of an individual SAG isolate cannot be explained by any one feature and is likely an emergent property of its combined functional capabilities and ecological context. SAG are emerging as important invasive pathobionts, not only in the head and neck but in other sterile cavities, and more research into this diverse group is required.

## 7 Data Availability Statement

Genome sequence data generated in this study have been deposited with the National Centre for Biotechnology Information (NCBI) under BioProject accession number PRJNA1528876. Raw Nanopore sequence read data have been deposited in the Sequence Read Archive (SRA) under the accession numbers SRR40721160 to SRR40721207 and the draft genome assemblies have been deposited under accession numbers SAMN63118671 to SAMN63118718 (**Supplementary Table 2**).

## Declaration of generative AI and AI-assisted technologies in the manuscript preparation process

During the preparation of this work, the authors used Claude (Anthropic; models Sonnet 5 and 5.5, Opus 5 and 5.5, and Fable 5.1) to assist with improving the language and clarity of the manuscript, and to assist with writing code and running certain bioinformatic analyses. The authors reviewed, tested, and edited all AI-assisted content and outputs as needed, and take full responsibility for the methodology, the analyses performed, and the content of the article.

